# Improved detection of gene fusions by applying statistical methods reveals new oncogenic RNA cancer drivers

**DOI:** 10.1101/659078

**Authors:** Roozbeh Dehghannasiri, Donald Eric Freeman, Milos Jordanski, Gillian L. Hsieh, Ana Damljanovic, Erik Lehnert, Julia Salzman

## Abstract

The extent to which gene fusions function as drivers of cancer remains a critical open question. Current algorithms do not sufficiently identify false-positive fusions arising during library preparation, sequencing, and alignment. Here, we introduce a new algorithm, DEEPEST, that uses statistical modeling to minimize false-positives while increasing the sensitivity of fusion detection. In 9,946 tumor RNA-sequencing datasets from The Cancer Genome Atlas (TCGA) across 33 tumor types, DEEPEST identifies 31,007 fusions, 30% more than identified by other methods, while calling ten-fold fewer false-positive fusions in non-transformed human tissues. We leverage the increased precision of DEEPEST to discover new cancer biology. For example, 888 new candidate oncogenes are identified based on over-representation in DEEPEST-Fusion calls, and 1,078 previously unreported fusions involving long intergenic noncoding RNAs partners, demonstrating a previously unappreciated prevalence and potential for function. Specific protein domains are enriched in DEEPEST calls, demonstrating a global selection for fusion functionality: kinase domains are nearly 2-fold more enriched in DEEPEST calls than expected by chance, as are domains involved in (anaerobic) metabolism and DNA binding. DEEPEST also reveals a high enrichment for fusions involving known and novel oncogenes in diseases including ovarian cancer, which has had minimal treatment advances in recent decades, finding that more than 50% of tumors harbor gene fusions predicted to be oncogenic. The statistical algorithms, population-level analytic framework, and the biological conclusions of DEEPEST call for increased attention to gene fusions as drivers of cancer and for future research into using fusions for targeted therapy.

**Significance:** Gene fusions are tumor-specific genomic aberrations and are among the most powerful biomarkers and drug targets in translational cancer biology. The advent of RNA-Seq technologies over the past decade has provided a unique opportunity for detecting novel fusions via deploying computational algorithms on public sequencing databases. Yet, precise fusion detection algorithms are still out of reach. We develop DEEPEST, a highly specific and efficient statistical pipeline specially designed for mining massive sequencing databases, and apply it to all 33 tumor types and 10,500 samples in The Cancer Genome Atlas database. We systematically profile the landscape of detected fusions via employing classic statistical models and identify several signatures of selection for fusions in tumors.

**Software availability:** DEEPEST-Fusion workflow with a detailed readme file is available as a Github repository: https://github.com/salzmanlab/DEEPEST-Fusion. In addition to the main workflow, which is based on CWL, example input and batch scripts (for job submission on local clusters), and codes for building the SBT files and SBT querying are provided in the repository. All custom scripts used for systematic analysis of fusions are also available in the same repository.

## Introduction

While genomic instability, resulting in DNA sequence composition that deviates from the reference, is a hallmark of human cancers, its functions have only partially been explained (Negrini et al., 2010). Point mutations and gene dosage effects result from genomic instability (Stranger et al., 2007), but they alone do not explain the origin of human cancers (Martincorena et al., 2015). Genomic instability also results in structural variation in DNA that creates rearrangements, including local duplications, deletions, inversions or larger scale intra- or inter-chromosomal rearrangements that can be processed into chimeric mRNAs, called gene fusions.

Gene fusions are known to drive some cancers and can be highly specific and personalized therapeutic targets; some of the most famous fusions are the BCR-ABL1 fusion in chronic myelogenous leukemia (CML), the EML4-ALK fusion in non-small lung cell carcinoma, TMPRSS2-ERG in prostate cancer, and FGFR3-TACC3 in a variety of cancers including glioblastoma multiforme (Nowell and Hungerford, 1960; Soda et al., 2007; Tomlins et al., 2008; Singh et al., 2012). Since fusions are generally absent in healthy tissues, they are among the most clinically relevant events in cancer to direct targeted therapy and to be used as effective diagnostic tools in early detection strategies using RNA or proteins; moreover, as they are truly specific to cancer, they have promising potential as neo-antigens (Zhang, Mardis and Maher, 2017; Ragonnaud and Holst, 2013; Liu and Mardis, 2017).

Because of this, clinicians and large sequencing consortia have made major efforts to identify fusions expressed in tumors via screening massive cancer sequencing datasets (Hu et al., 2017; Alaei-Mahabadi et al., 2016; Gao et al., 2018; Stransky et al., 2014; Yoshihara et al., 2014). However, these attempts are limited by critical roadblocks: current algorithms suffer from high false positive rates and unknown false negative rates. Thus, ad hoc choices have been made in calling and analyzing fusions including taking the consensus of multiple algorithms and filtering lists of fusions using manual approaches (Wang et al., 2015; Abate et al., 2012; Liu et al., 2015). These approaches lead to what third-party reviews agree is imprecise fusion discovery and bias against discovering novel oncogenes (Liu et al., 2015; Carrara et al., 2013; Kumar et al., 2016). This suboptimal performance becomes more problematic when fusion detection is deployed on large cancer sequencing datasets that contain thousands or tens of thousands of samples. In such scenarios, precise fusion detection must overcome a problem of multiple hypothesis testing: each algorithm is testing for fusions thousands of times, a regime known to introduce false positives. To overcome these problems, the field has turned to consensus-based approaches, where multiple algorithms are run in parallel (Gao et al., 2018), and a meta-caller allows ‘voting’ to produce the final list of fusions. This is also unsatisfactory, as it introduces false negatives.

Both shortcomings in the ascertainment of fusions by existing algorithms and using recurrence alone to assess fusions’ function have limited the use of fusions to discover new cancer biology. As one of many examples, a recent study of more than 400 pancreatic cancers found no recurrent gene fusions, raising the question whether this is due to high false negative rates or whether this means that fusions are not drivers in the disease (Bailey et al., 2016). Recurrence of fusions is currently one of the only standards in the field used to assess the functionality of fusions, but the most frequently expressed fusions may not be the most carcinogenic (Saramäki et al., 2008); on the other hand, there may still be many undiscovered gene fusions that drive cancer.

Thus, the critical question “are gene fusions under-appreciated drivers of cancer?” is still unanswered. In this paper, we provide several contributions that more precisely define this question and provide important advances to answering it. First, we provide a new algorithm that has significant advance in precision for unbiased fusion detection at exon boundaries in massive genomics datasets. Our new algorithm, Data-Enriched Efficient PrEcise STatistical Fusion detection (DEEPEST-Fusion), is a second-generation fusion algorithm with significant computational and algorithmic advance over our previously developed MACHETE algorithm (Hsieh et al., 2017). A key innovation in DEEPEST-Fusion is its statistical test of fusion prevalence across population, which can identify false positives in a global unbiased manner.

The precision and efficient implementation of DEEPEST-Fusion allowed us to conduct an unbiased screen for expressed fusions occurring at annotated exon boundaries (based on GRCh38) in a cohort of 10,521 RNA-sequencing datasets, including 9,946 tumor samples and 575 normal (tumor adjacent) samples, across the entire 33 tumor types of The Cancer Genome Atlas (TCGA). Beyond recovery of known fusions, DEEPEST-Fusion discovers novel fusions and new cancer biology.

While frequent recurrence of gene fusions has been considered a hallmark of a selective event during tumor initiation, and this recurrence has historically been the only evidence available to support that a fusion drives a cancer, private or very rare gene fusions are beginning to be considered potential functional drivers (Latysheva and Babu, 2016). However, the high false positive rates in published algorithms prevent a statistical analysis of whether reported private or rare gene fusions exhibit a signature of selection across massive tumor transcriptome databases, such as TCGA. For the first time, we have formulated statistical tests for non-neutral selection of fusion expression by calculating the expected rates of rarely recurrent gene fusions and partner genes, enrichment of gene families such as kinase genes or those curated in Catalogue Of Somatic Mutations In Cancer (COSMIC) (Forbes et al., 2014), and enrichment for protein domains or pairs of protein domains present exclusively in fusions. These analyses reveal a significant signal for selection of gene fusions. The statistical tests provide a basis for identifying new candidate oncogenes and driver and druggable fusions.

To illustrate one of our findings, a large fraction of ovarian serous cystadenocarcinoma cancers has until now lacked explanatory drivers beyond nearly universal TP53 mutations and defects in homologous recombination pathways. Because TP53 mutations create genome instability, a testable hypothesis is that TP53 mutations permit the development of rare or private driver fusions in ovarian cancers, and the fusions have been missed due to biases in currently available algorithms. We apply DEEPEST-Fusion to RNA-Seq data from bulk tumors and find that 94.6% of the ovarian tumors we screened have detectable fusions, half of the ovarian cancer tumors express gene fusions involving a known COSMIC gene, and 36% have fusions involving genes in a kinase pathway.

In summary, DEEPEST-Fusion is an advance in accuracy for fusion detection in massive RNA-Seq data sets. The algorithm is reproducible, publicly available, and can be easily run in a dockerized container (Methods). Its results have important biological implications: DEEPEST-Fusion, applied in conjunction with new statistical analysis to the entire TCGA database, reveals a signature of fusion expression consistent with the existence of under-appreciated drivers of human cancer, including selection for rare or private gene fusions with implications from basic biology to the clinic.

## Results

### DEEPEST-Fusion is a new statistical algorithm for gene fusion discovery in massive public databases

We engineered a new statistical algorithm, Data-Enriched Efficient PrEcise STatistical Fusion detection (DEEPEST-Fusion), to discover and estimate the prevalence of gene fusions in massive numbers of data sets. Here we have applied DEEPEST-Fusion to ∼10K datasets, but in principle, DEEPEST can be applied to 100K, 1 million or more samples. DEEPEST-Fusion includes key innovations such as controlling false positives arising from analysis of massive RNA-Seq data sets for fusion discovery, a problem conceptually analogous to multiple hypothesis testing via p-values, which cannot be solved by direct application of common FDR controlling procedures which rely on the assumption of a uniform distribution of p-values under the null hypothesis.

The DEEPEST-Fusion pipeline contains two main computational steps: (1) junction nomination component which is run on a subset of all samples to be analyzed called “the discovery set” and (2) statistical testing of nominated junctions on all analyzed samples, “the test set”. In this paper, we have used all samples as the discovery set, but this set could be a fraction of RNA-seq data if desired.

Step 1 includes running KNIFE (Szabo et al., 2015) to detect chimeric junctions, defined as a splicing event between two distinct genes, whose exons are on the same chromosome and within the distance of 1Mb, and a method based on the MACHETE algorithm to detect chimeric junctions with partner exons being farther than 1Mb from each other or on different chromosomes/strands (Fig. 1). Putative fusions are nominated from the initial database by employing a null statistical model of read alignment profiles that models the effect of junction sequence composition and gene abundance in generating false positive fusions (Supplemental File and Methods). This step relies on new computational engineering, which restructures the MACHETE pipeline into an efficient reproducible publicly available workflow based on dockerized containers, using the Common Workflow Language (CWL). Another advance in DEEPEST-Fusion over MACHETE is further improvement of sensitivity by including gold standard cancer fusions in the junction nomination step of MACHETE, which makes DEEPEST-Fusion easily portable to clinical settings where clinicians desire precise identification of a set of known fusions. For this purpose, we used fusions curated in ChimerDB 3.0 (Lee et al., 2016).

**Figure 1:**
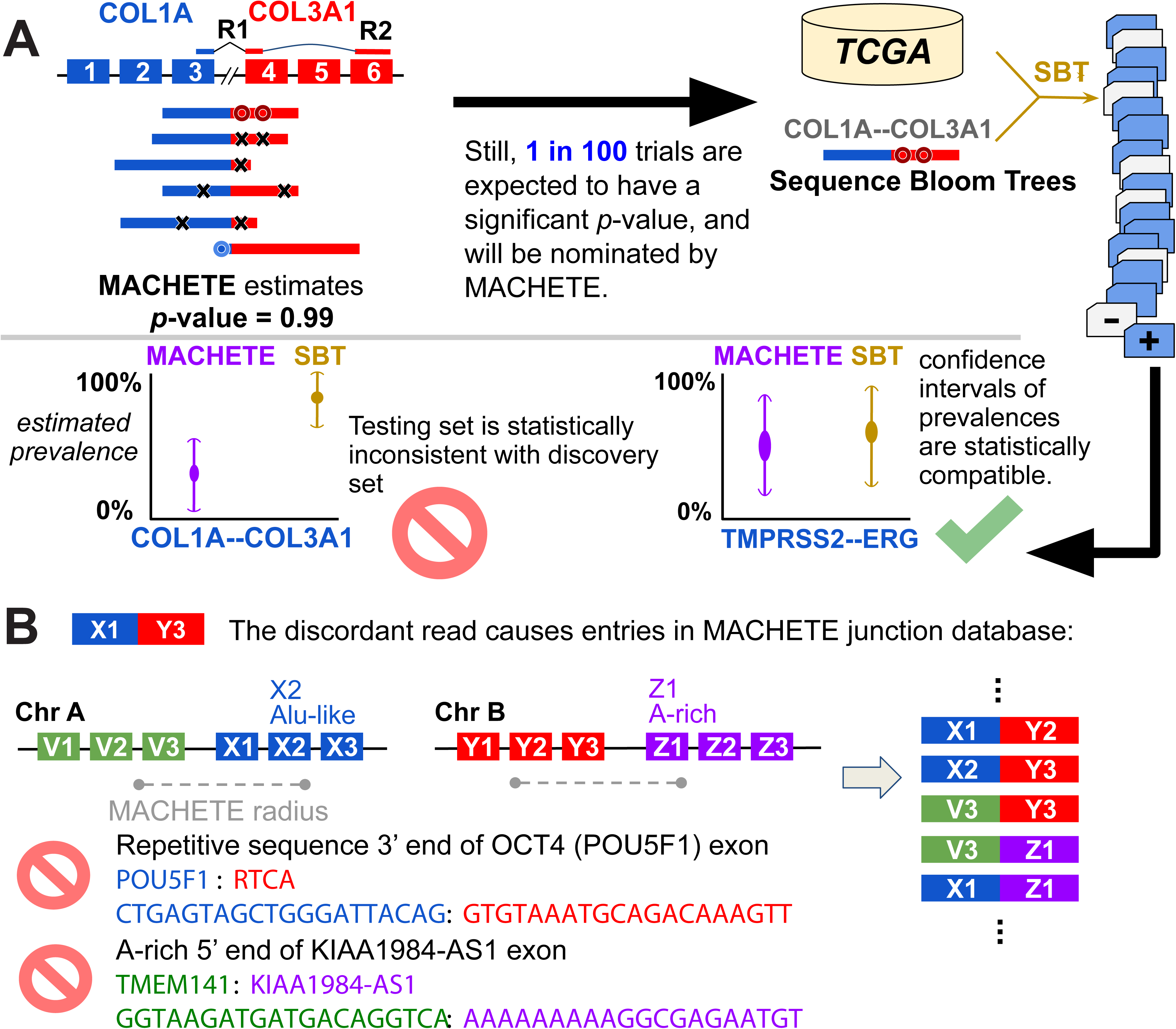
Origin and identification of FP from running DEEPEST on 1000s of samples. (a) DEEPEST uses all reads, including those censored by other algorithms, to generate an empirical p-values for each candidate fusion. Sequence Bloom Trees (SBTs), together with further statistical modeling are used to identify FP arising from testing on multiple samples (The first black arrow shows the motivation for designing the SBT step). (b) cDNA or mapping artifacts result in the inclusion of exon-exon junctions from all combinations of exons within a fixed genomic radius of X1 with all exons in the radius of Y3. Some such exon junctions will include degenerate sequences which cannot be mapped uniquely, and thus DEEPEST blinds itself to detection of fusions containing such highly degenerate sequences (for example, due to Alu exonization) or with polyA stretches at the 5’ end.

In Step 2, the statistical refinement step, DEEPEST-Fusion uses new statistical approaches, based on orthogonal sequence level queries using a sequence bloom tree (SBT; Solomon and Kingsford, 2016), a method that indexes the sequence composition of genomic data sets and can rapidly query whether specific kmers appear in the corpus. This step is modular and can in principle be applied to any fusion discovery algorithm to identify false positives resulting from multiple testing, a major challenge brought on by running discovery algorithms on massive datasets. Fusions nominated by the junction nomination component are subjected to a second statistical test: they are efficiently tested in the discovery set along with an arbitrarily large number of added samples in the test set, here tens of thousands of samples, by rapid queries using SBT. This step further decreases the false positive identification of fusions beyond MACHETE, which has been already shown to have higher specificity than any other published algorithm (Hsieh et al., 2017). Intuitively, this step checks whether the prevalence of fusions found by running MACHETE (or KNIFE) is statistically consistent with the estimated prevalence using a string-query based approach (such as SBT). Since the SBT has perfect sensitivity by searching merely by looking at fusion-junctional sequences, samples could be positive for a fusion by SBT yet negative by MACHETE, which requires discordant reads to nominate fusions (Hsieh et al., 2017). For a fusion to be called by DEEPEST-Fusion, it should have an SBT detection frequency that is statistically consistent with the estimated prevalence by the junction nomination component and additionally pass statistical filtering such as a test for repetitive sequences near exon boundaries (Figure 1 and Supplemental File).

DEEPEST-Fusion does not require human guidance, is fully automated, and can be applied to any paired-end RNA-Seq database by leveraging the massive computational power of cloud platforms. A web-based user-friendly version of the pipeline has been implemented on the Seven Bridges Cancer Genomics Cloud (http://www.cancergenomicscloud.org/), which allows a user to run the workflow either by uploading RNA-seq data or using RNA databases already available on CGC (Currently, the average cost of running the workflow for a single TCGA sample on the cloud is roughly $3). Moreover, most parts are portable as they are dockerized and can be easily exported to many platforms using a description given by the Common Workflow Language (the workflow codes are provided in a Github repository). In addition, the custom scripts and files needed for systematic analysis of fusions are provided in the Github repository.

### DEEPEST-Fusion improves sensitivity and specificity of fusion detection

We first evaluated DEEPEST-Fusion false positive and false negative rates on fusion positive benchmarking sample data used by third parties to assess the performance of 14 state-of-the-art algorithms (Hass et al., 2017). On each dataset, DEEPEST-Fusion has 100% positive predictive value (PPV), the ratio of the number of true positive calls to the total number of calls, higher than all 14 other state-of-the-art algorithms (Supplemental Figure 1), DEEPEST-Fusion has a comparable, though numerically higher PPV than the next best algorithms (PRADA). For this analysis, we only applied the first component of DEEPEST-Fusion, which is based on MACHETE, as the SBT refinement step utilizes the statistical power across a large cohort of samples, which is not the case for simulated datasets.

Because simulations can only model errors with known sources, it is common for algorithms to perform differently on real and simulated data; for example, simulated data does not model reverse transcriptase template switching, or chimeras arising from ligation or PCR artifacts. Thus, in addition to evaluating the performance of DEEPEST on simulated data, we performed a thorough computational study of DEEPEST’s performance on real data. To further evaluate the false positive rate of DEEPEST-Fusion, we applied it to several hundred normal datasets, including GTEx (Lonsdale et al., 2013) and TCGA normal samples. Notably, DEEPEST-Fusion calls 80% fewer fusions in GTEx samples than does STAR-Fusion (Supplemental Figure 2A), an algorithm used in a recent pan-cancer TCGA analysis (Gao et al., 2018). In addition, DEEPEST-Fusion reports fewer fusions (509 fusions) on TCGA normal samples compared to the 3,128 calls in the same samples by TumorFusions (Hu et al., 2017) (Supplemental Figure 2B), which is a TCGA fusion list based on PRADA (Torres-García et al., 2014). This provides evidence that, unlike other algorithms, DEEPEST-Fusion retains the specificity seen in simulations in real tissue samples.

We ran DEEPEST-Fusion on the entire TCGA RNA-seq corpus: 9,946 tumor samples across all 33 tumor types. DEEPEST-Fusion detects 31,007 fusions across the entire TCGA corpus. Consistent with what is known about tumor-type-specific gene fusion expression, DEEPEST-Fusion reports the highest abundance of fusions in sarcoma (SARC), uterine corpus endometrial carcinoma (UCEC), and esophageal carcinoma (ESCA) tumor types, and the fewest number of detected fusions in thyroid carcinoma (THCA), testicular germ cell tumors (TGCT), and (uveal melanoma) UVM (for the rest of paper, we use TCGA tumor type abbreviations).

While calling significantly fewer fusions in normal samples, DEEPEST-Fusion identifies significantly more fusions in TCGA tumor samples compared to recent surveys of the same samples (Gao et al., 2018 and Hu et al., 2017), the former is based on STAR-Fusion that is more sensitive in simulated data. While some fusion algorithms might exhibit better sensitivity (at the cost of higher false positive rates) on simulated datasets, DEEPEST-Fusion is more sensitive in real cancer datasets (Supplemental Figure 2 C,D). When samples shared between three lists are considered, DEEPEST-Fusion detects much more fusions (29,820 fusions, compared to 23,624 fusions in (Gao et al., 2018) and 19,846 fusions by TumorFusions) and substantially fewer calls in real normal datasets (Supplemental Figure 2A,B), suggesting that the modeling employed by DEEPEST-Fusion is a better fit for real data. DEEPEST-Fusion-only fusions are enriched in cancers known to have high genomic instability (ESCA, OV, STAD, SARC) compared to fusions found only by TumorFusions and (Gao et al., 2018) (Supplemental Figure 2D). Together, this implies that DEEPEST-Fusion is more specific on simulated and real data and identifies more high confidence fusions on real data.

Because fusions between exons that are closer to each other than 1MB in the reference and transcribed on the same strand could be due to local DNA variation or transcriptional or post-transcriptional splicing, for example into circRNA (Salzman et al., 2013), we define an “extreme fusion” to be a fusion that joins exons that are farther than 1MB apart, are on opposite strands, or are on different chromosomes and profile the distribution of DEEPEST-Fusion-called fusions as a function of extreme characteristics. Around 24% of fusions have both partner genes transcribed from the same chromosome and strand and within 1MB, 22% are on the same chromosome and strand but separated by at least 1MB, 23% are strand crosses with genes being on opposite strands, and 31% are interchromosomal fusions (Figure 2A).

**Figure 2:**
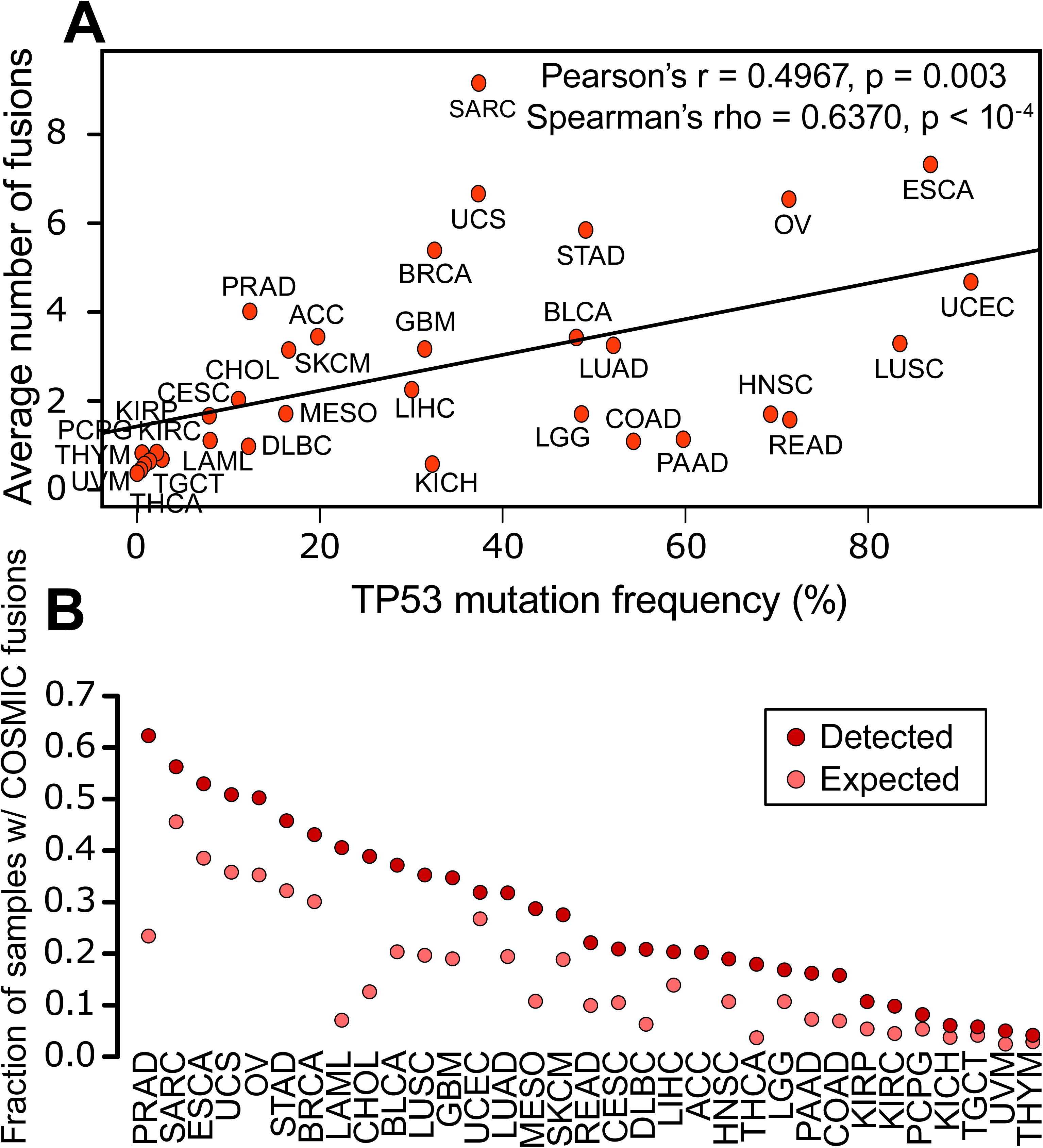
Association of fusions with genomic instability. (A) DEEPEST fusions are significantly correlated with the TP53 mutation frequency. (B) Detected fusions are highly enriched in genes catalogued by COSMIC. For all tumor types except for UCS, UCEC, PCPG, UVM, TGCT, KICH, THYM, and ACC, the observed fraction significantly exceeds the expected fraction based on the null hypothesis of random pairing (Bonferroni corrected FDR < 0.05).

DEEPEST-Fusion finds 1,486 recurrent fusions (512 distinct recurrent fusions), called in at least two tumors within a tumor type (Figure 2B). Restricted to a single tumor type, most fusions have low levels of recurrent gene fusions (Fig. 2C) with exceptions of the well-known TMPRSS2-ERG in PRAD (182 samples, 36.3% of tumor samples), PML-RARA in LAML (14 samples, 8% of tumor samples), and DHRS2-GSTM4 in BLCA (Figure 2C). Many gene fusions are detected in diverse cancers, for example, MRPS16-CFAP70 and FGFR3-TACC3 (10 cancer types) (Figure 2D).

Around 41% of DEEPEST-Fusion’s 31,007 fusions (12,196 fusions) are new TCGA fusions in the sense that they are only found by DEEPEST-Fusion on TCGA datasets (Supplemental Figure 2C). Far fewer fusions are found only by one of the other algorithms (4,402 fusions in TumorFusions and 5,860 fusions in (Gao et al., 2018)) (Supplemental Figure 2C). We further investigated DEEPEST-Fusion-only fusions and queried them through FusionHub portal (https://fusionhub.persistent.co.in/) to see if they are present in any other fusion database and found that 9,272 distinct fusions (i.e., gene pairs) were not present in any other fusion database (Supplemental Table 1). Included in this list are 157 previously unreported recurrent fusions (Supplemental Table 1, Supplemental Figure 3), including a novel recurrent fusion for PRAD involving SCHLAP1, a long noncoding RNA known to have driving oncogenic activities in the prostate cancer (Prensner et al., 2013).

### Statistical analysis of fusion prevalence in large data queries improves precision of calls

The SBT refinement step is a critical innovation of the DEEPEST-Fusion pipeline and substantially improved its specificity by removing fusion calls likely to be false positives. Such fusions passed the first component of DEEPEST due to multiple hypothesis testing but were flagged by the statistical SBT refinement step (Supplemental Figure 4A).

The main power of the novel refinement step is in its unbiasedness: it is agnostic to gene name or ontology (such as pseudogenes, paralogs, synonymous genes, and duplicated genes) used by conventional filters, and which can lead to true positives being removed from lists (eg RUNX1--RUNX1T1). Furthermore, most fusions removed during the SBT refinement step would have passed such filters (Supplemental Figure 4B).

To further evaluate the performance of the SBT-based refinement component, we extracted two groups of fusions from the fusions called by the junction nomination component: (1) likely false positives (LFP) that are fusions with SBT hits in the GTEx samples and (2) likely true positives (LTP) that contain fusions shared between the first component, (Gao et al., 2018), and TumorFusions. Note that LFP fusions are defined on the basis of GTEx data, whereas the SBT refinement step does not “touch” GTEx data, meaning that the LFP fusion set can be used as a test set for the specificity of the SBT refinement step. A fusion is removed by the SBT refinement step if its two-sided binomial p-value is less than 0.05, an arbitrary and conventional statistical threshold. To evaluate the precision of the SBT step, we treated each two-sided p-value as a continuous measure and stratified the p-values for LFP and LTP sets (Supplemental Figure 4C). The p-values of LFP fusions (with GTEx SBT hits) are significantly smaller than those of LTP fusions (Mann-Whitney U test, p-value < 2.2e-16), implying that the majority of them are filtered out by the SBT statistical refinement step. On the other hand, LTPs possess higher two-sided p-values, which indicates that they would pass the refinement step. In summary, while the statistical refinement step improves DEEPEST-Fusion calls, the approach outlined is also a general methodology for increasing the precision of RNA variant calls on massive datasets.

### DEEPEST-Fusion identifies novel long noncoding RNA fusions

A major category of fusions identified by DEEPEST-Fusion are fusions that involve long noncoding RNAs (LncRNAs), which have been overlooked by other methods due to bioinformatic or heuristic filters. Fusions involving well-studied (and thus named) long noncoding RNAs have been appreciated to have oncogenic potential (Latysheva and Babu, 2016; Chunru and Yang, 2018; Huarte, 2015). Due to their functions in regulating cellular homeostasis, fusions that involve LncRNA have the potential to contribute or drive oncogenic phenotypes (Huarte, 2015; Kopp and Mendel, 2018). Genome-wide pan-cancer analyses have previously not profiled those with the LINC annotation. We found that fusions involving long noncoding RNAs (as annotated by the Ensembl database release 89) are abundant in tumors (10% of fusion calls involve LncRNAs and 20% of tumors are found to have at least one fusion involving an LncRNA; Supplemental Table 3). A large fraction of LncRNA fusions (∼30%, or 994 fusions) involves LINC RNAs, which have been overlooked by previous methods due to their biases and heuristic filters for discarding “uncharacterized” genes.

### Prevalence of fusions found by DEEPEST is correlated with genome instability

To more carefully study DEEPEST-Fusion precision in primary tumors where no ground truth is known, we use the well-studied and generally cytogenetically simple acute myeloid leukemia (LAML) as the best approximation for ground truth. In a large cohort of LAML samples investigated through both next-generation sequencing and cytogenetics by a large consortium (Cancer Genome Atlas Research Network, 2013; Papaemmanuil et al., 2016), DEEPEST-Fusion improves the rate of true positive recovery by reporting 96 validated fusions. In total, DEEPEST-Fusion detects 3,530 fusions of the 8,523 fusions reported in TCGA marker papers for 17 different tumor types (Supplemental Table 1).

As another computational test of whether DEEPEST-Fusion maintains high precision in a variety of solid tumors that have more complex cytogenetics than LAML, we tested whether the abundance of detected fusions per cancer is correlated with the mutation rate of TP53, which is an orthogonal measure of a tumor’s genome instability (Forment et al., 2012). The average number of fusions per sample identified by DEEPEST-Fusion has a higher (and significant) correlation with TP53 mutation rate across tumor types compared to (Gao et al., 2018) and TumorFusions (Pearson correlation 0.497, 0.38, and 0.31 respectively; Spearman’s correlation 0.637, 0.596, and 0.54 respectively; Figure 3A). To our knowledge for the first time, we found that there is a significant correlation between fusion abundance and TP53 mutation frequency as two orthogonal measures of genomic instability. DEEPEST-Fusion calls more fusions in tumors with high TP53 mutation rates in less cytogenetically complex tumors, while retaining tight control of false positives in other samples.

**Figure 3:**
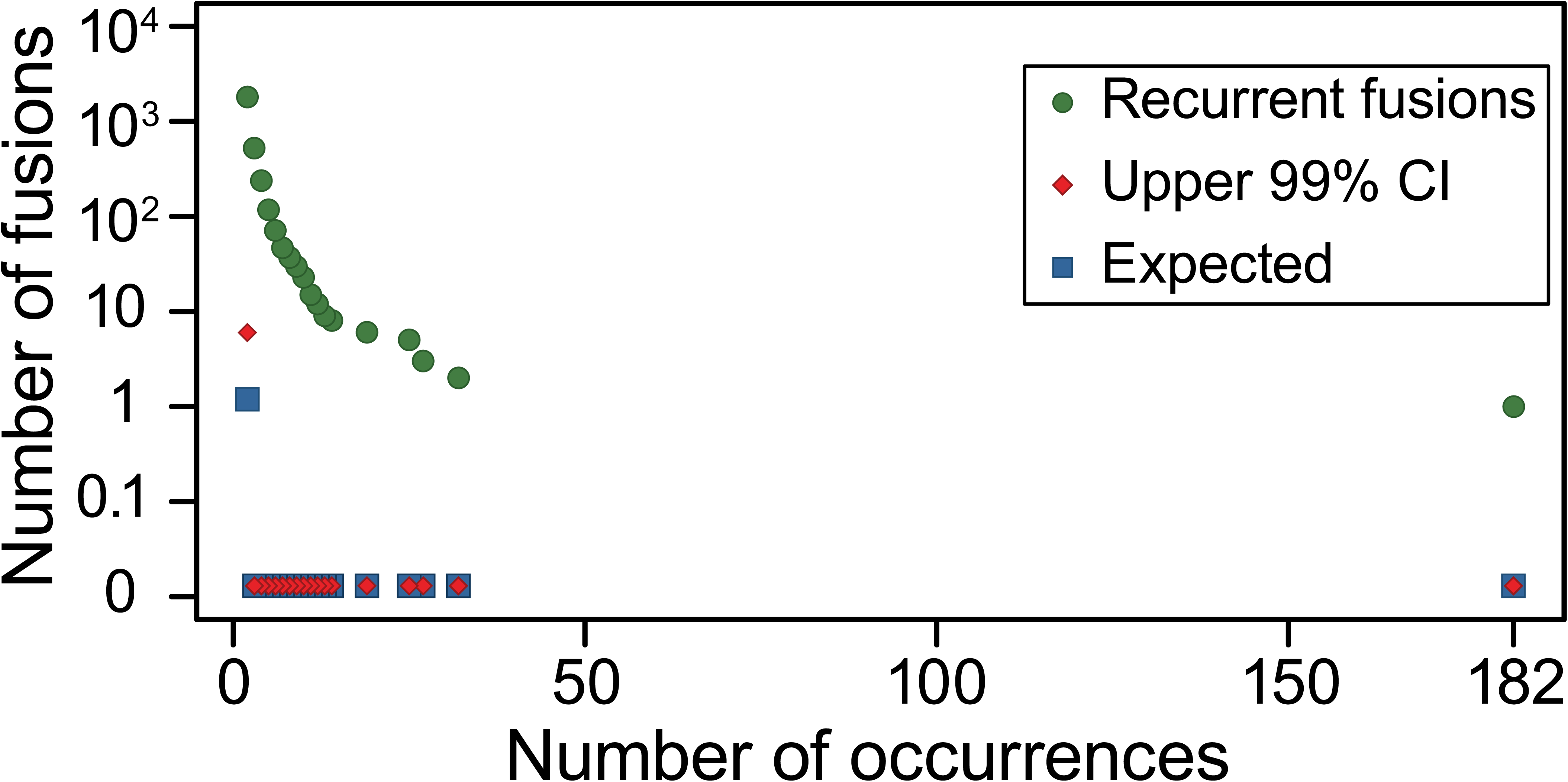
Statistical analysis of recurrent fusions. Observed number of recurrent fusions that occur more than x times is significantly higher than the expectation and the upper 99\% confidence interval expected in the null (Benjamini-Yekutieli FDR control at level 0.01).

### DEEPEST-Fusion identifies a positive selection on fusions containing known oncogenes

Gene fusions could arise from random reassortment of DNA sequences due to deficiencies in the structural integrity of DNA in tumors with no functional impact, or they can be driving events, such as for BCR-ABL1 in Chronic Myelogenous Leukemia (Nowell and Hungerford, 1960). In our first global computational tests of whether fusions are passengers or drivers, we tested whether DEEPEST-Fusion-called fusions are enriched in genes known to play oncogenic roles. Since functional gene ontologies are not used by DEEPEST-Fusion, this analysis provides an independent test of whether fusions are enriched for genes in known cancer pathways. For each tumor type, we tested for enrichment of the 719 genes present in the COSMIC Cancer Gene Census database (Forbes et al., 2014).

Under the null hypothesis of random pairing of genes in fusions, we compared the observed to expected fraction of samples containing at least one fusion with a COSMIC gene by conditioning on the total number of fusions detected for a tumor type, where tumor types with more detected fusions are expected to have a higher ratio of samples with COSMIC fusions (Methods). The largest enrichment for COSMIC genes is in PRAD (3 fold change vs expected fraction, p-value < 1e-6), THCA (4.9 fold change vs expected fraction, p-value < 1e-6), and LAML (5.6 fold change vs expected fraction, p-value < 1e-6) (Figure 3B, Supplemental Table 4). This is expected because the most frequent gene fusions in PRAD involve the ETS family of transcription factors (which are cataloged as COSMIC genes), THCA tumors are highly enriched for kinase fusions, and LAML is a disease where fusions, including known drivers, have been intensively studied and therefore their partners are annotated as COSMIC genes. Most tumor types lack prevalent recurrent gene fusions, and thus there is no a priori bias that fusions will be enriched for COSMIC genes in other tumor types.

In PRAD, SARC, ESCA, UCS, and OV, the fraction of samples with fusions containing a COSMIC partner exceeds 50%, a rate much greater than expected by chance, the null fraction of samples with COSMIC fusions is 45% for SARC and less than 40% for other tumor types (Figure 3B, and Supplemental Table 4). In more than 90.7% (Bonferroni corrected FDR <0.05) of the tumor samples we studied, COSMIC genes are statistically enriched above the background rate. Together this is strong evidence for a positive selection pressure on gene fusions in various tumor types, including cancers such as OV where fusions are currently not considered to play a driving role.

### Statistical analysis of rare fusions shows a selection for fusions in more than 11% of TCGA tumors

Fusion recurrence is considered to be evidence that a fusion plays a driving role. This argument grew out of work focused on point mutations in cancer genomes (Rowley, 2001). However, the total number of possible gene fusions (the sample space) greatly exceeds the sample space of point mutations. The number of potential gene fusions scales quadratically with the number of genes in the genome (in the samples we analyzed ∼22,000 genes were expressed). This means that there are up to 625 million potential gene fusions, more than an order of magnitude greater than the number of possible point mutations which is bounded by the number of protein-coding bases in the transcribed genome (∼30*10^6^). Therefore, fusions could be strongly selected for in tumors without observing high levels of recurrence.

If a moderate fraction of human genes could function as oncogenes when participating in fusions, rare fusion expression is expected in a population-level survey, even one as large as the TCGA cohort.

To account for this effect, we formalized a statistical test for whether the prevalence of rare recurrent fusions fits a model of neutral selection by a null distribution where fusion expression arises by chance, the theory of which was worked out in (Henze, 1998, Methods). We mapped the probability of observing recurrent gene fusions to a familiar problem in statistics: if k balls (corresponding to the number of observed fusions) are thrown into n boxes (corresponding to the total number of possible gene pairs), how many boxes are expected to have c or more balls? In other words, given the number of detected fusions, how many of them are expected to be called for at least c samples?

The most prevalent fusions expected under neutral selection would be observed only 2 times, and we would expect to observe only 5 such fusions (Figure 4) making this and thousands of other fusions highly unlikely to be observed under the null hypothesis. Controlling for multiple hypothesis testing, this analysis recovers several known recurrent fusions including TMPRSS2-ERG, PML-RARA, and FGFR3-TACC3 (Kakizuka et al., 1991; Singh et al., 2012), The second most prevalent gene fusion is DHRS2-GSTM4 (Figure 2), which involves the tumor suppressor gene DHRS2 and the oncogene GSTM4 (Luo et al., 2009).

**Figure 4:**
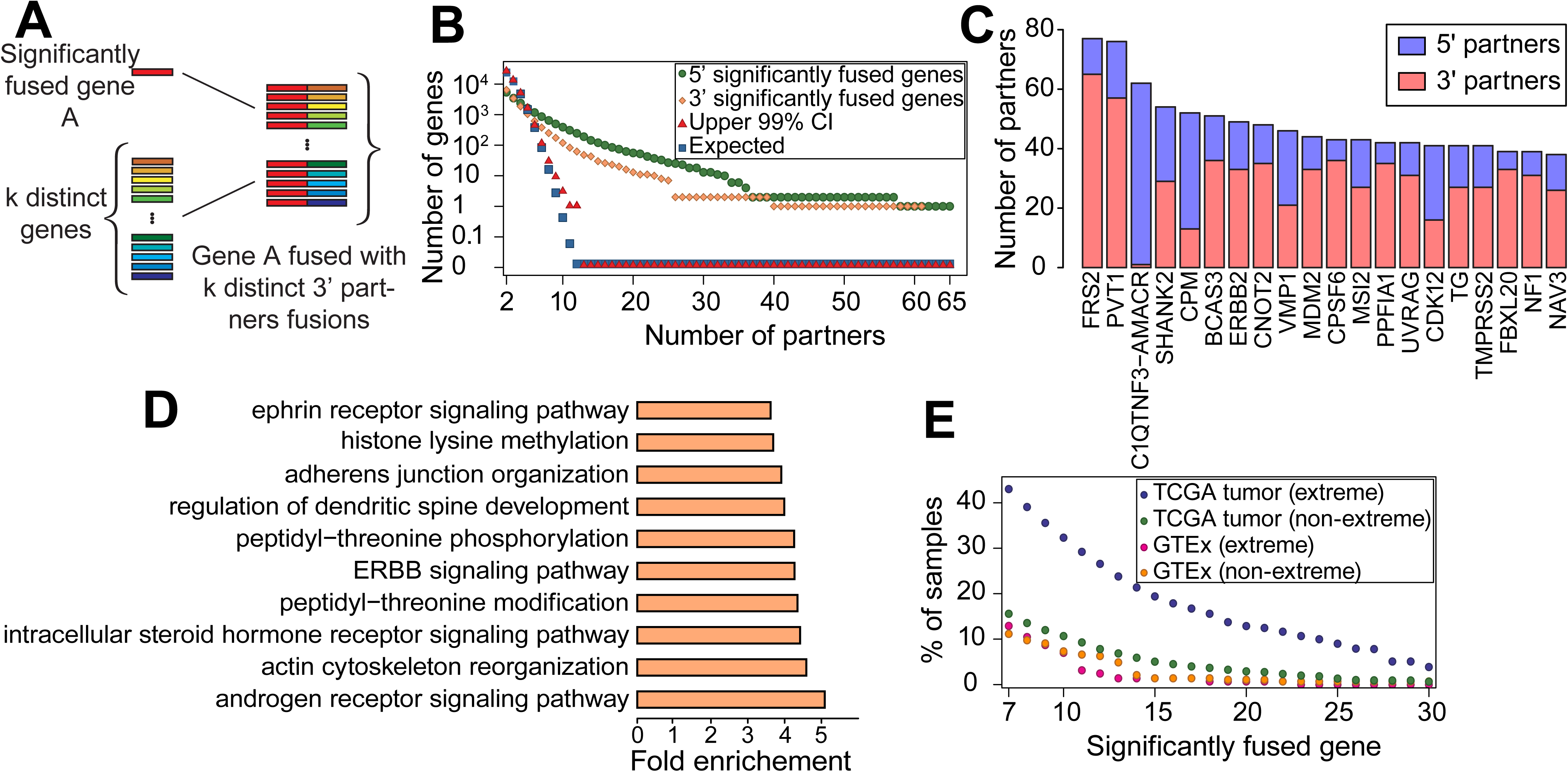
(A) Significantly fused genes: those paired with multiple partners (counted by counting distinct fusion pairs) (B) Significantly fused genes are observed at rates higher than expected by chance; genes with k > 10 partners would be unlikely to occur by random assortment of genes into fusions (FDR corrected p-value < 0.01). (C) Genes with the highest number of distinct gene partners (D) GO enrichment analysis of significantly fused genes shows enrichment in pathways known to regulate cancer growth. (E) Tumors are highly enriched for extreme fusions involving significantly fused genes.

This analysis reveals new evidence that recurrent fusions are selected for in diverse tumors; RPS6KB1-VMP1, a fusion between the ribosomal protein kinase (Cai et al., 2015) and a vacuolar protein (VMP1) present in a 8 tumor types, is the most prevalent detected gene fusion, after TMPRSS2-ERG, across the entire TCGA cohort and supports findings by previous studies that these fusions have a driving role (Inaki et al., 2011, Blum et al., 2016; Figure 2C). Globally, 14% of the fusions (1,486) found by DEEPEST-Fusion are observed at higher rates than expected by chance (p < 1e-6); more than 11.9% of tumors (1,181) have recurrent fusions (Figure 4 and Supplemental Table 1).

### Recurrently fused genes distinguish tumors from non-neoplastic tissue and are fused in more than 30% of TCGA tumors

If many genes could serve as oncogenic fusion partners, fusions under selection could be private, yet partners could be much more prevalent than would be expected by chance. To test whether 3’ or 5’ partner genes are over-represented in fusions found by DEEPEST-Fusion in the TCGA cohort, we used the “balls in boxes” null distribution above where boxes correspond to all possible 3’ (respectively 5’) partners (expressed genes) and balls correspond to the total number of fusion pairs (i.e., 31,007 fusions) detected across all samples. We map the coincidence of c balls in one box to c distinct 5’ (resp. 3’) partner genes being paired with one 3’ (resp. 5’) partner and call genes with statistically significant numbers of 5’ and 3’ partners “significantly fused” (Figure 5A and B).

**Figure 5:**
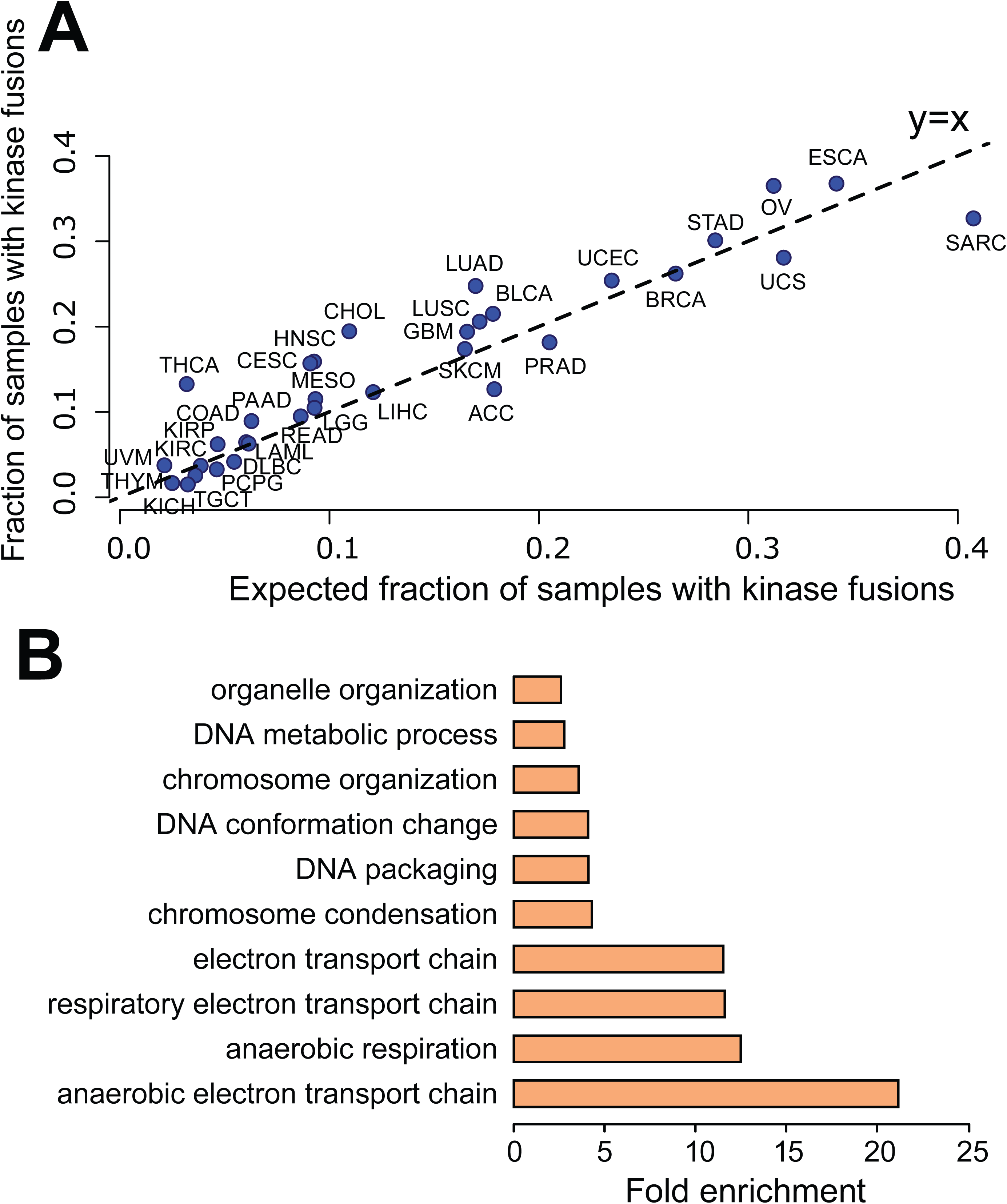
Protein domain analysis. (A) Analysis of the fraction of samples containing kinase fusions reveals that THCA, CHOL, LUAD and OV, and many other tumor types, have significant high enrichment of kinase fusions in addition of high overall rates (B) GO analysis identifies enrichment of cellular metabolism and DNA organization in the protein domains enriched in all fusion transcripts.

The number of significantly-fused 5’ and 3’ partners is large: DEEPEST-Fusion reports 864 recurrent 5’ partners and 378 recurrent 3’ partners, both having p-values < 1e-5 (Figure 5B), when only 110 genes with more than 6 partners would be expected by chance (Supplemental Table 2); 190 and 48 genes are found in fusions as significantly fused 5’ and 3’ partner genes with more than 12 partners, respectively, when no such genes would be expected by chance. The most significant 5’ partner gene is FRS2, a docking protein that is critical in FGF receptor signaling (Hadari et al., 2001); FRS2 fusions are detected in 52 tumors or in 0.5% of TCGA cases. Other highly significant recurrent partners include PVT1, ERBB2 (HER2), known oncogenes, and tumor suppressors such as MDM2, which negatively regulates P53 (Kubbutat et al., 1997) and UVRAG (Liang et al., 2006) (Figure 5C). The most promiscuous 3’ partner genes are CPM, a gene regulating innate immune development (39 partners), and the gene C1QTNF3-AMACR (61 partners) (Figure 5C). Other genes with the highest numbers of distinct 5’ partners include CDK12, a cyclin-dependent kinase emerging as a target in cancer therapy (Lui et al., 2018), and well-known tumor suppressors such as RAD51B (Thacker, 2005). We also found 31 noncoding RNAs as significantly fused genes. PVT1, noncoding RNAs of unknown function: AC134511.1, AC025165.3, and LINC00511 have the most 3’ partners; and BCAR4, PVT1, and noncoding RNAs of unknown function: AP005135.1 and AC020637.1 have the most 5’ partners (Supplemental Table 2). While some of these noncoding RNAs such as PVT1 (Guan et al., 2007), LINC00511 (Sun et al., 2016), and BCAR4 (Xing et al., 2014) have been shown to act as oncogenes, our findings both re-identify them and call for further investigation into the potential driving roles of other significantly-fused noncoding RNAs.

While the list of significantly fused genes includes well-known cancer genes as described above, only 15.5% of them (193 genes) are currently annotated as COSMIC genes, i.e., 888 significantly fused genes are new candidate oncogenes, calling for further functional investigation of these genes. This is an enrichment for COSMIC annotation, but is not exhaustive: significantly fused genes are a large new class of potential oncogenes and tumor suppressors that function through gene rearrangement rather than gain or loss of function through point mutation. To functionally annotate significantly-fused partner genes, we carried out GO enrichment analysis. The GO biological processes that had highest enrichment and significance (Binomial test, Bonferroni-corrected FDR < 0.05) coincided with major cancer pathways such as androgen signaling pathway, ERBB signaling pathway, and ephrin receptor signaling pathway (Figure 5D, Supplemental Table 2).

To further support the role of significantly fused genes in cancer, we evaluated the rate that such genes are detected in TCGA tumor and GTEx samples as a function of (a) the number of partners and (b) the nature of the rearrangement underlying the fusion: “extreme” events that bring together two exons that are farther than 1MB apart, on opposite strands, or on different chromosomes in the reference, and all others (“non-extreme”) (Figure 5E). Non-extreme events could arise through small scale genomic duplication or transcriptional readthrough coupled to “backsplicing” to generate circRNA (Szabo et al., 2015; Hsieh et al., 2017). Globally, DEEPEST-Fusion detects fusions including significantly fused genes (>10 partners) at a much higher rate in TCGA tumors (7,050 fusions in ∼34% of samples) than in GTEx controls (29 such fusions in ∼9% of samples), despite GTEx samples being sequenced at an average depth of 50 million reads, roughly similar depth to tumor samples.

The deviation between the fraction of such fusions in TCGA versus GTEx increases with the number of partners of the significantly fused gene such that among those with at least 23 partners, only two fusions are detected in GTEx (0.7% of samples), while 1,845 such fusions are detected in 1,202 TCGA tumors (12.1% of samples). Notably, the two fusions detected in GTEx are PVT1-MYC and FRS2-CPSF6, both fusions are ‘non-extreme’, splicing detected between two genes transcribed in the same orientation with promoters < 200kb from each other, events which could arise from somatic or germline variation or transcriptional readthrough.

Controlling for differing numbers of samples in tumor and non-neoplastic cohorts, significantly fused genes are enriched in TCGA tumors compared to GTEx: there is a ∼4 fold enrichment for significantly fused genes (>10 partners) in extreme fusions in TCGA versus GTEx and these fusions are the dominant contributor to significantly fused genes (Figure 5E). This implies a selection for expression of such fusions by tumors. Further, fusions involving significantly fused genes in TCGA and GTEx samples have distinguishing structural features. The large majority of such fusions in tumors arise from extreme rearrangements (Supplemental Figure 6) regardless of the number of partners a significantly fused gene contains. More than 90% of fusions in TCGA that involve significantly fused genes with at least 23 partners are extreme, whereas no such GTEx fusions are extreme (Supplemental Figure 6). This again implies a tumor-specific selection for extreme fusions, which increase the complexity of partners available to significantly fused genes. Together, analysis of rare recurrent gene fusions and recurrent 3’ and 5’ partners identify hundreds of new genes, which constitute a significant fraction of gene fusions and is underlined by selection on such fusions that grows as a function of the number of partners used by the significantly fused gene.

DEEPEST-Fusion finds higher enrichment of significantly-fused genes in larger number of tumors compared to other TCGA fusions lists (Supplemental Figure 7), calling significantly-fused genes with >10 partners in ∼50%; and with >20 partners in ∼70% more samples compared to recent studies (>10 partners: DEEPEST: 3,705, (Gao et al., 2018): 2,570,TumorFusions: 2,787 samples; >20 partners: DEEPEST: 1,479, (Gao et al., 2018): 823,TumorFusions: 958 samples). While 7.6% of DEEPEST fusions have a gene with >20 partners, only 4.8% and 5.6% of fusions in (Gao et al., 2018) and TumorFusions, respectively have such genes. While FRS2 is found to have the highest number of partners in all three lists, DEEPEST identifies 65 3’ partners, which is larger than other two lists: 41 and 52 partners by (Gao et al., 2018) and TumorFusions, respectively.

### Lung Adenocarcinoma and Serous Ovarian Carcinoma have high statistical enrichment for kinase fusions

The most common genetic lesions in OV and LUAD cancers is TP53 mutation, present in 85.8% of OV and 52.12% of LUAD cases (cBioPortal, retrieved November 19, 2018, Cerami et al., 2013), although there is a debate in the literature that this prevalence is an underestimate. However, TP53 mutations are not sufficient to cause cancers (Martincorena et al., 2015). In OV, the explanatory driving events are as yet unknown (Bowtell et al., 2015). We tested the hypothesis that genome instability in OV could generate fusions responsible for driving some fraction of these cancers, which might have been missed because of shortcomings in fusion detection sensitivity. The rate of kinase fusions is statistically significantly higher than would be expected by chance, supporting a selection for and driving role of kinase fusions in these tumor types. DEEPEST-Fusion predicts that 37% of ovarian tumors (Binomial test, p-value < 1e-5) and 25% of lung adenocarcinoma tumors (Binomial test, p-value < 1e-5) contain kinase fusions (Figure 6A, Supplemental Table 4), a rate higher than what would be expected based on the null assumption of random pairing of genes in fusions. Other cancers with high enrichment of kinase fusions include: THCA (13.3% of samples, p-value < 1e-6), HNSC (16% of samples, p-value < 1e-6) and CESC (15.7% of samples, p-value = 7.7e-5) (Figure 6A, Supplemental Table 4).

### Positive selection for fusions to rewire the cancer proteome

To test if there is selection on the protein domains included in fusions, we compared the rate at which each protein domain occurs in the reference proteome to its prevalence in the DEEPEST-Fusion-called fusion proteome. This analysis identified a set of 120 domains that are statistically enriched in fusion proteins. The most highly enriched domains are AT_hook, a DNA binding motif found, for example in the SWI/SNF complex, NUP84_NUP100, a domain present in some nucleoporins, and Per1 a domain involved in lipid remodeling, all present at 15 times higher frequency than the reference proteome (p-value << 1e-10) (Supplemental Table 3). Tyrosine kinase domains are 1.8 fold enriched in fusions compared to the reference proteome (p << 1e-10). To functionally characterize the 120 domains enriched in fusions proteins, we performed Gene Ontology (GO) enrichment analysis using dcGOR (Fang, 2014) and identified over-represented biological processes among these domains (Binomial test, Benjamini-Yekutieli FDR control, corrected FDR < 0.05): the enriched domains were involved in (anaerobic) electron transport, chromosome condensation and organization, and DNA metabolism or organization (Figure 6B, Supplemental Table 3).

To find the set of domain-pairs enriched in fusions, we compared the observed frequency of each domain pair against the null probability of random pairing between domains. 226 domain pairs are enriched above background (Bonferroni corrected FDR < 0.05), among the highest enriched domain pairs are NHR2--RUNT, RUNT--TAFH and RUNT--zf-MYND in the in-frame fusion protein RUNX1--RUNX1T1 detected in LAML samples (Supplemental Table 3).

Because enrichment of protein domain pairs could be sensitive to how we model the null distribution, we formulated a test for selection of fusion proteins containing two in-frame domains where the ‘most pessimistic’ null distribution for our problem can be computed in closed-form. This analysis considers only fusions whose 5’ and 3’ parent genes contain only one annotated domain. Out of 3,388 fusions with one-domain parental genes, 681 fusions with 2 domains were observed, whereas only 282 were expected by chance under a closed-form, conservative null distribution (p-value < 1e-5) (Supplemental File), strong evidence for selection of such fusions that couple intact domains in the fusion protein.

In addition to the above enrichment, 17% of all DEEPEST-Fusion fusions result in proteins that have protein domain-pairs that do not exist in the reference proteome. These pairs include well-known driving fusions such as the domain pairs Pkinase_Tyr-TACC and I-set-TACC in FGFR3-TACC3, but also include 9,500 other domain-pairs not found in the reference proteome, which implies their potential for tumor-specific function (Supplemental Table 3).

## Discussion

Some of the first oncogenes were discovered with statistical modeling that linked inherited mutations and cancer risk (Knudson, 1971). The advent of high-throughput sequencing has promised the discovery of novel oncogenes, which can inform basic biology and provide therapeutic targets or biomarkers (Cibulskis et al., 2013; Lawrence et al., 2014). However, unbiased, sequencing-based methodologies for the discovery of novel oncogenic gene fusions have been only partially successful.

DEEPEST-Fusion is a unified, reproducible statistical algorithm to detect gene fusions in large-scale RNA-Seq datasets without human-guided filtering. DEEPEST-Fusion has significantly lower false positive rates than other algorithms. The unguided DEEPEST-Fusion filters have not sacrificed detection of known true positives. Further, DEEPEST-Fusion assigns a statistical score that can be used to prioritize fusions on the basis of statistical support, rather than the absolute read counts supporting the fusion. Such a statistical score is unavailable in other algorithms but of potential scientific and clinical utility as the discovery rate and the trade-off between sensitivity and specificity of DEEPEST can be tuned by modifying the threshold on scoring.

Although many likely driving and druggable gene fusions have been identified by high-throughput sequencing, studies reporting them have either a non-tested or non-trivial false positive rate even using heuristic or ontological filters, making those fusions unreliable for clinical use. Similar problems also limit the sensitivity in screens of massive data sets to discover fusions, novel oncogenes or signatures of evolutionary advantage for rare or private gene fusions. To illustrate DEEPEST-Fusion potential clinical contribution, we integrated DEEPEST-Fusion-detected fusions with OncoKB (Chakravarty et al., 2017), which is a recent curated database of precision oncology with drugs stratified by evidence that they interact with specific proteins and found druggable fusions in 327 tumors (3.3%) (Supplemental Figure 8). However, the current list of potentially druggable fusions is biased towards genes that have classically been considered oncogenes, and mainly kinases. By increasing the precision of fusion calls, the analysis in this paper opens the door for further functional and clinical studies to better identify drug targets and repurpose already validated drugs that could target fusions.

The DEEPEST-Fusion algorithm improves detection of gene fusions that have been missed by other algorithms’ lists of “high confidence” gene fusions. Analysis of these gene fusions uncovers new cancer biology. First, we find evidence that gene fusions are more prevalent than previously thought in a variety of tumors including high grade serous ovarian cancers. Second, our computational analysis suggests the hypothesis that gene fusions involving kinases and perhaps other genes are a contributing driver of these cancers. Further, fusions in ovarian and most tumor types are under selection to include gene families that are known to drive cancers, such as kinases and genes annotated as COSMIC.

DEEPEST-Fusion allows for rigorous and unbiased quantification of gene fusions at annotated exonic boundaries and for tests of whether partners in gene fusions that may be rare or private are present at greater frequencies than would be expected due to chance. The results in this paper establish statistical evidence that gene fusions and the partner genes involved in fusions are under a much greater selective pressure than previously appreciated: under a highly stringent definition of an enriched partner, more than 10-20% of all TCGA tumors profiled harbor a gene fusion including a gene under selection by tumors to be involved in gene fusions that require large scale genomic rearrangement. Future work profiling normal cohorts will distinguish whether fusions including these genes that found in GTEx arise due to variation at the DNA sequence, transcriptional or post-transcriptional levels. Significantly, this discovery required new statistical analysis of rare gene fusions, using large numbers of samples to increase power to detect selection for gene fusion expression by tumors.

Together, the results in this paper lead to a new model that fusions may be lesions like point mutations, present across tumors rather than tumor-defining, and suggests that by focusing on one tumor type to detect recurrence, and by relying on classical metrics for recurrent and selection, some important cancer biology is lost. Further, the computational evidence in this paper suggests rare fusions are drivers of a substantial fraction of tumors.

While DEEPEST-Fusion has increased the accuracy of fusion detection, there are obvious extensions of this work. Above, we described fusions between genes where one gene can be drugged by existing therapies, including ERBB2 (HER2). Further work with a clinical focus is needed to determine the extent of potentially druggable fusions identified by DEEPEST-Fusion. Second, we have limited our analysis to fusion RNAs that occur at annotated exon-exon boundaries; we believe that extending the statistical approaches used to discover gene fusions may allow us to relax the requirement that gene fusions be detected at annotated exonic sequences, without sacrificing the false positive rate. Doing so will provide a more powerful test of whether genomic instability in cancers results in gene fusions that are a “passenger” of this instability or that have currently under-appreciated functional and perhaps clinical importance by increasing detection of fusions containing cryptic exons and perhaps the signal for significantly fused partners.

## Methods

### An enhanced statistical fusion detection framework for large scale genomics

We used the discovery set to generate a list of fusions passing MACHETE statistical bar (Figure 1). We then queried all data sets for any fusions found in any discovery set (Figure 1) and estimated the incidence of each fusion with SBTs (Solomon and Kingsford, 2016). Next, we used standard binomial confidence intervals to test for consistency of the rate that fusions were present in the samples used in MACHETE discovery step and the rate that they were found in the SBT. Fusion sequences that were more prevalent across the entire data set that is statistically compatible with the predicted prevalence from the discovery set were excluded from the final list of fusions (Figure 1).

For intuition on why this step is important, consider the scheme in Figure 1: given an exon-exon junction query sequence that could be generated by sequencing errors convolved with gene homology or ligation artifacts, SBTs will not consider the alignment profile of all reads aligning to this junction as MACHETE does, e.g., reads with errors or evidence of other artifacts, because reads with mismatches with the query sequence are by definition censored by the SBT. As a result, the SBT, like other algorithms, can have a high false positive rate due to: (a) false positives intrinsic to the Bloom filters used in the SBT; (b) false positive identification of putative fusions due to events such as the one depicted in Figure 1, even in the presence of a null false positive rate by the SBT itself. False positives as in (b) can arise as follows: if a single artifact (e.g. a ligation artifact between two highly expressed genes) in a single sample passes MACHETE statistical threshold in the discovery step, this artifact will be included as a query sequence, and the SBT could detect it at a high frequency because the statistical models employed by MACHETE are not used by the SBT (Figure 1). Testing for the consistency of the rate of each sequence being detected in the discovery set with its prevalence as estimated by SBTs controls for the multiple testing bias described above (Figure 1). Technical details of the statistical framework, post-processing of DEEPEST-Fusion output files, and generation of SBT queries are described in the Supplemental File.

We built SBT filters for all TCGA samples across various TCGA projects using SBT default parameters (kmer index size 20 and a minimum count of 3 for adding a kmer to a bloom filter). The 40mer flanking the fusion junction (20 nucleotides on the 5’ side and 20 nucleotides on the 3’ side) is retrieved for each fusion nominated by the junction nomination component and then the TCGA cancers are queried for the fusion query sequences using the SBT filters built for them. We used a more stringent value of 0.9 for the sensitivity threshold (instead of its default value 0.8) to improve the specificity of DEEPEST-Fusion; this parameter could be changed by other users to tune specificity. For each TCGA cancer type, a fasta file containing the 40-nucleotide sequences for all fusions called by the first component is queried using the SBT filters built for that type. Then for each fusion junction, its detection frequencies by the first component and SBT query are compared and if they are statistically consistent, the fusion would pass the SBT refinement step; otherwise, it would be discarded.

### Sequence Bloom Tree Methodology

Sequence Bloom Trees (SBTs, Solomon and Kingsford, 2016) are data structures developed to quickly query many files of data of short-read sequences from RNA-Seq data (and other data) for a particular sequence. These structures build on the concept of Bloom filters. The authors published software, which was subsequently Dockerized and wrapped in the Common Workflow Language (CWL) for use on the Seven Bridges Cancer Genomics Cloud (Lau et al.; 2017). The supplemental file contains technical details about the methodology used.

### Null probability for recurrent fusions

For g genes, there are g(g-1) possible fusions. If n fusions are detected by DEEPEST-Fusion across all tumors, let X be the number of recurrent fusions. In our analysis, there are g = 22,000 different gene names in DEEPEST-Fusion report files and we have reported n = 31,007 fusions. The probability that no fusion is recurrent can be computed using the Poisson approximation for the birthday coincidences problem with lambda= n(n-1)/(2g(g-1)): P(X=0)∼= e^-lambda=0.451. As shown in Figure 4, the expected value of the number of recurrent fusions (with a frequency of at most 2) is 5.

### Calculations for the expected number of recurrent 5’ and 3’ partners

As a test of the likelihood of observing our results, we employ a statistical model of the probability of observing as many or more recurrent 5’ and 3’ partners under the assumption that the genes in each fusion pair are randomly chosen from all expressed genes. To do this, we employ the generalized birthday model from (Henze, 1998):

First, we consider all g = 22,000 expressed genes as boxes, which represent each potential 5’ partner gene. Next, we consider the distribution of the number of distinct 3’ partners for each gene if 3’ partner genes, which we take to be numbered balls, were thrown at random into boxes. When the first ball arrives in the box j_1, this represents that the first observed fusion on our list has gene j_1 on its 5’ side. At the end of this process, we have thrown n = 31,007 fusions (balls) into n boxes. For a given c, the number of balls occupying a single box, we can calculate the probability of having X_{g,c} boxes with at least c balls. The distribution of X_{g,c} has been shown to be a Poisson distribution Po(t^c/c!) ((Henze, 1998; Theorem 2.2), where t = n/(g^{1-1/c}). We perform the following calculations to find significant recurrent 5’ gene partners. For each c, we find the expected number and the 99% upper confidence interval of the number of boxes (5’ genes) that have at least c balls (distinct 3’ partners) according to the null distribution. For statistical analysis, a significance level of 0.01 was considered. Moreover, since we are testing multiple hypotheses in our analysis, we adopt the Benjamini-Hochberg-Yekutieli FDR control procedure (Benjamini and Yekutieli, 2001) and correct the significance value for each c. For each c, we construct the confidence interval at level (1 - corrected significance level) (Figure 5B). Similarly, we can find the expectation and upper confidence interval (after Benjamini-Hochberg-Yekutieli correction) for each number of recurrent 3’ genes that have at least c 5’ gene partners (Figure 5B).

We provide a table of p-values for each observation of the number of recurrent genes with greater than or equal to c partners in Figure 5 by the formula 1-F(number of 3’ (5’) partners with at least c 5’ (3’) partners), where F is the cumulative distribution function of the Poisson distribution Po(t^c/c!) (Supplemental Table 2).

### Description of tumor types analyzed in this study

In this pan-cancer study, we have processed 9,946 tumor and 575 matched-normal TCGA samples across 33 different cancer types (Supplemental Figure 5): adrenocortical carcinoma (ACC), bladder urothelial carcinoma (BCLA), breast invasive carcinoma (BRCA), cervical squamous cell carcinoma and endocervical adenocarcinoma (CESC), cholangiocarcinoma (CHOL), colon adenocarcinoma (COAD), lymphoid neoplasm diffuse large B-cell lymphoma (DLBC), esophageal carcinoma (ESCA), glioblastoma multiforme (GBM), head and neck squamous cell carcinoma (HNSC), kidney chromophobe (KICH), kidney renal clear cell carcinoma (KIRC), kidney renal papillary cell carcinoma (KIRP), acute myeloid leukemia (AML), brain lower grade glioma (LGG), liver hepatocellular carcinoma (LIHC), lung adenocarcinoma (LUAD), lung squamous cell carcinoma (LUSC), mesothelioma (MESO), ovarian serous cystadenocarcinoma (OV), pancreatic adenocarcinoma (PAAD), pheochromocytoma and paraganglioma (PCPG), prostate adenocarcinoma (PRAD), rectum adenocarcinoma (READ), sarcoma (SARC), skin cutaneous melanoma (SKCM), stomach adenocarcinoma (STAD), testicular germ cell tumors (TGCT), thyroid carcinoma (THCA), thymoma (THYM), uterine corpus endometrial carcinoma (UCEC), uterine carcinosarcoma (UCS), uveal melanoma (UVM). Raw paired-end fastq files for the TCGA RNA-Seq samples were accessed via the Data Browser tool provided in the Seven Bridges Cancer Genomics Cloud (CGC) (https://cgc.sbgenomics.com/datasets/#/tcga/data-browser). The matched normal samples span the following tumor types: BLCA, BRCA, CESC, CHOL, COAD, ESCA, GBM, HNSC, KICH, LUAD, LUSC, PAAD, PCPG, PRAD, READ, SARC, SKCM, STAD, THCA, THYM, and UCEC. GTEx samples for three different tissues: ovary (109 samples), blood (50 samples), and cerebral cortex (128 samples), were downloaded via the Database of Genotypes and Phenotypes (dbGaP) controlled access protocols and NCBI SRA Run Selector (https://www.ncbi.nlm.nih.gov/Traces/study/?acc=SRP012682)

### File downloads

- List of COSMIC genes (downloaded on 9/28/2018): http://cancer.sanger.ac.uk/cosmic/download
- List of kinase genes (Manning et al., 2002, downloaded on 10/11/2018): http://kinase.com/static/colt/data/human/kinome/tables/Kincat_Hsap.08.02.xls
- ChimerDB files (Lee et al., 2016, downloaded on 12/11/2016) http://203.255.191.229:8080/chimerdbv31/mdownload.cdb
- Mutation rates of the TP53 locus found in each available cancer subtype of the Cancer Genome Atlas database were accessed through the cBioPortal cancer genomics portal (http://www.cbioportal.org), on September 21, 2018.
- Protein domain information for linear transcriptome was downloaded by accessing the Ensembl BioMart search tool version 89 (http://may2017.archive.ensembl.org/biomart/martview) on 11/2/2018.
- List of duplicate genes (downloaded on 9/7/2018): http://dgd.genouest.org/listRegion/raw/homo_sapiens/all%3A0..x/
- List of gene synonym names (downloaded on 4/24/2018): ftp://ftp.ncbi.nih.gov/gene/DATA/GENE_INFO/Mammalia/Homo_sapiens.gene_info.gz
- List of pseudogenes was retrieved via the GeneCards human gene database portal (https://www.genecards.org) search tool on 9/8/2018.
- List of human paralogous gene pairs was retrieved by the **Ensembl BioMart** search tool (http://www.ensembl.org/biomart/martview/) on 9/8/2018, by selecting Ensembl Genes 94 and Human genes (GRCh38.p12) as the dataset, and Homologues as the Attributes.

## Acknowledgments

All RNA-Seq data was generated by The Cancer Genome Atlas project funded by the NCI and NHGRI. Information about TCGA and the investigators and institutions that constitute the TCGA Research Network can be found at https://cancergenome.nih.gov. We thank Steven Artandi, Rajat Rohatgi, and Peter Wang for useful discussions and suggestions on the DEEPEST-Fusion algorithm; Nathan Watson for kind help with code implementation on the CGC; Robert Bierman, Elisabeth Meyer, Julia Olivieri, and members of the Salzman lab for feedback on the manuscript. This work was supported by NIGMS grant [R01 GM116847], a JIMB seed grant, an NSF CAREER Award [MCB-1552196], McCormick-Gabilan Fellowship, and a Baxter Family Fellowship to J.S.. J.S. is an Alfred P. Sloan fellow in Computational & Evolutionary Molecular Biology. R.D. is supported by the Cancer Systems Biology Scholars program at Stanford [R25 CA180993]. The Seven Bridges NCI Cancer Genomics Cloud pilot and work by EL were supported in part by the funds from the National Cancer Institute, National Institutes of Health, Department of Health and Human Services, under Contract No. HHSN261201400008C. This research benefited from the use of credits from the National Institutes of Health (NIH) Cloud Credits Model Pilot, a component of the NIH Big Data to Knowledge (BD2K) program.

## Competing Interests

None.

## List of Figures

**Supplemental Figure 1:**
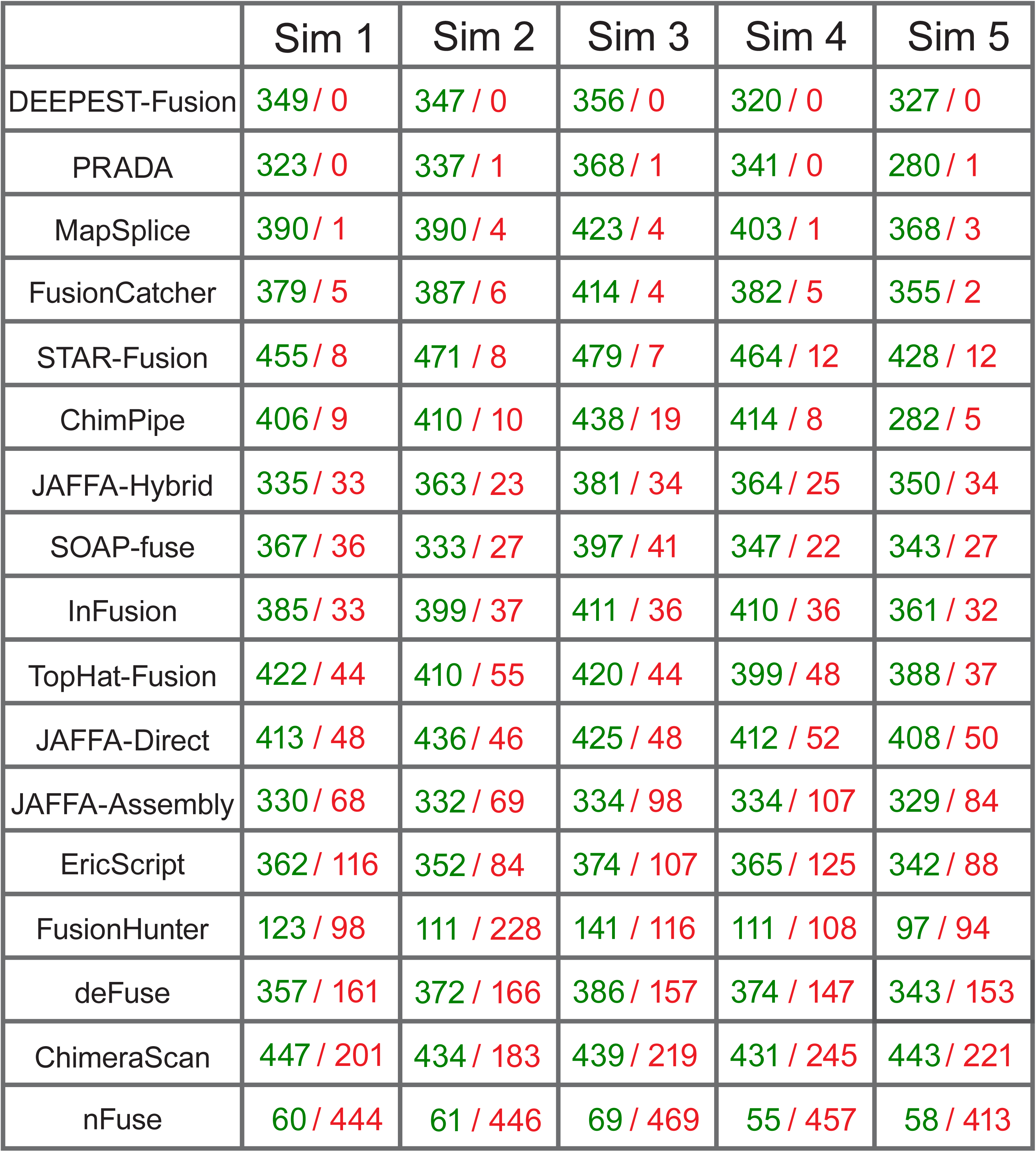
Comparison of DEEPEST (the first component based on MACHETE) with 14 other fusion detection methods based on the simulated datasets. In this comparison, we applied only the first component of DEEPEST. DEEPEST calls no FPs across all five datasets and maintains an acceptable sensitivity rate, while other methods suffer from high false positive rates. Each simulated dataset (Sim 1 up to Sim 5) has 500 true positive fusions. The first number (in green) and the second number (in red) are the number of TPs and FPs called by the method, respectively. The datasets were downloaded from: https://github.com/STAR-Fusion/STAR-Fusion_benchmarking_data.

**Supplemental Figure 2:**
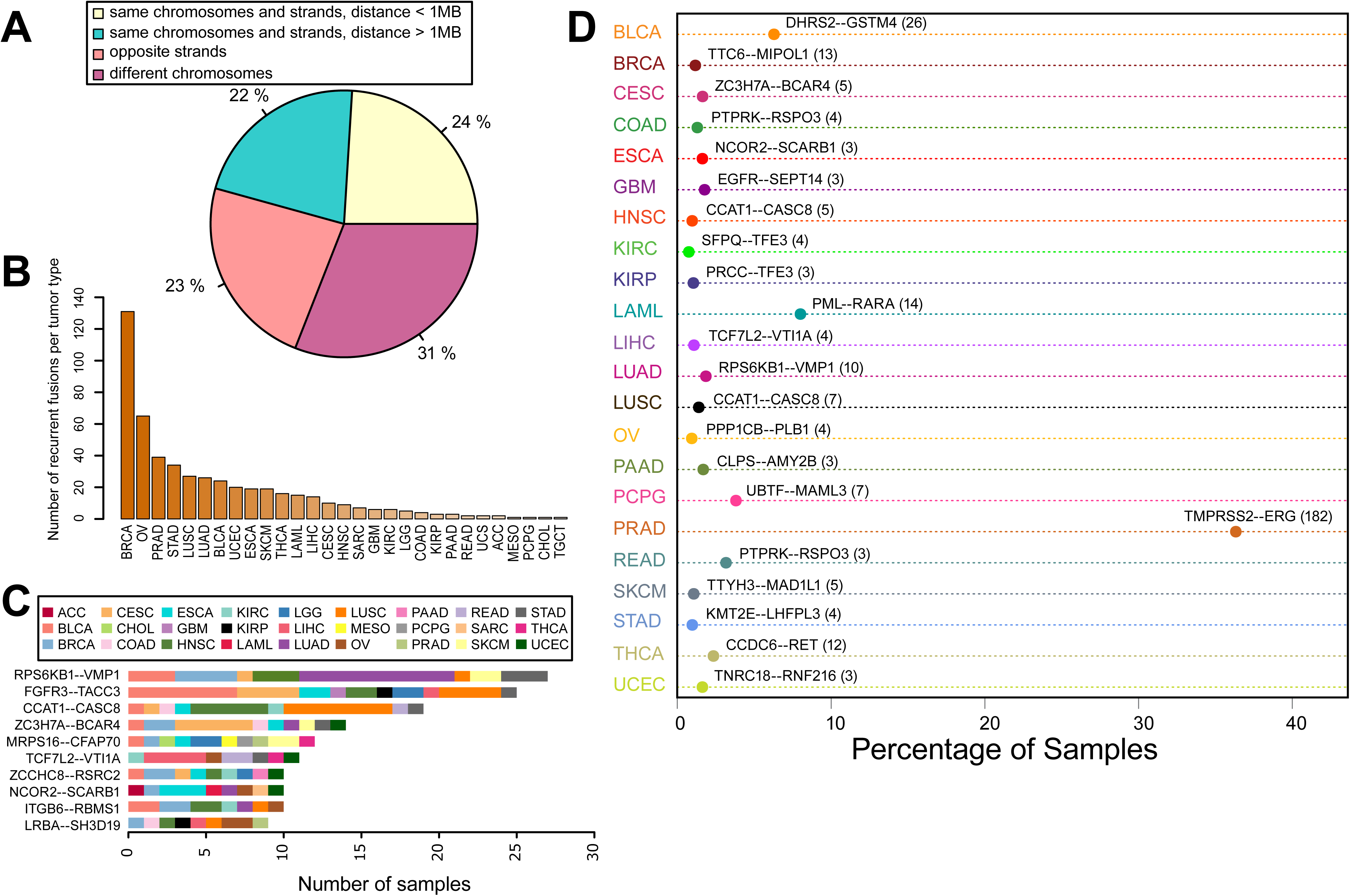
The landscape of detected fusions. (A) The relative position of the partner exons in the detected fusions. (B) The number of recurrent fusions for each tumor type. (C) The most recurrent fusion for each tumor type. (D) Fusions with the most diverse tumor types.

**Supplemental Figure 3:**
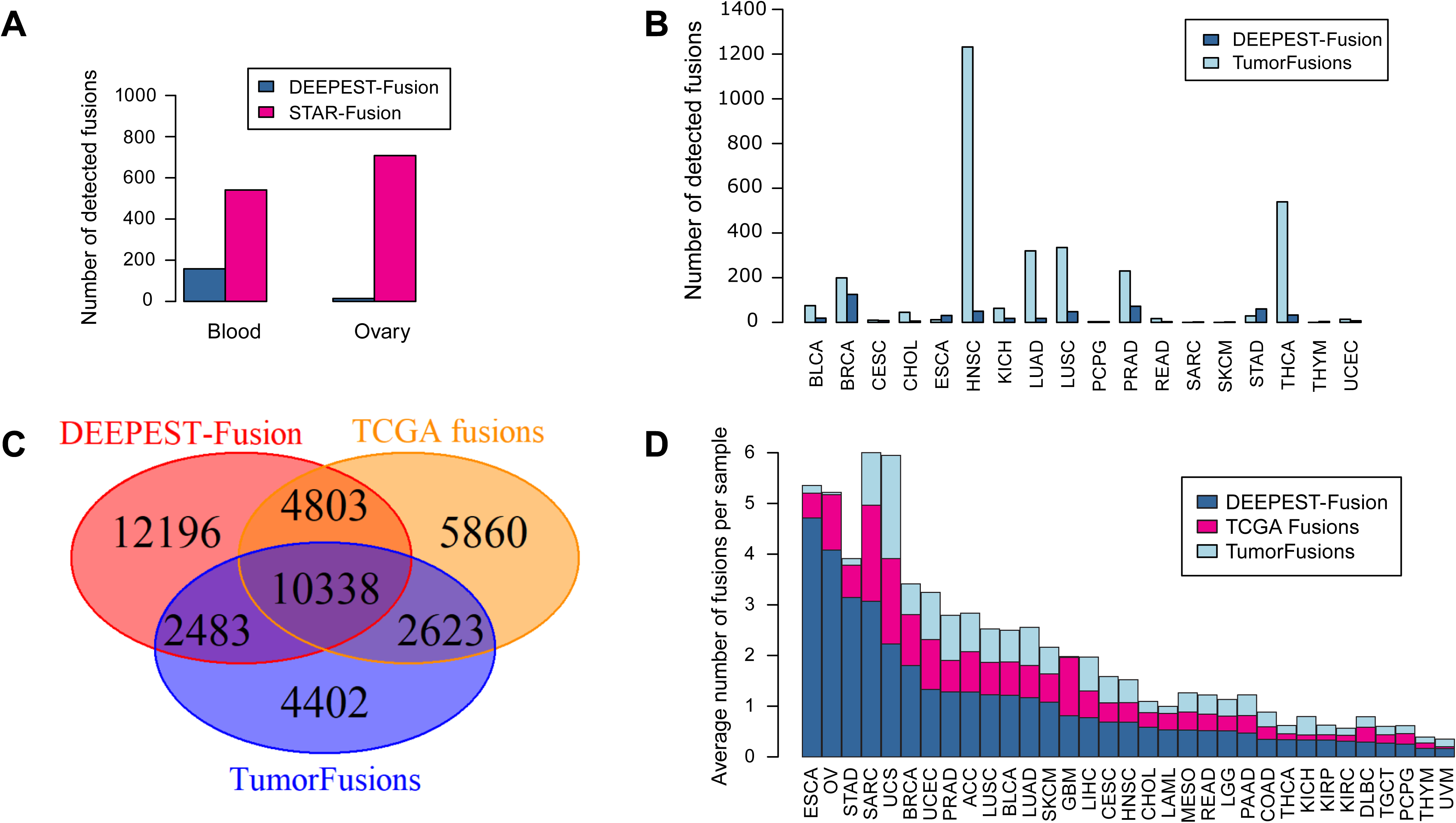
Benchmarking of DEEPEST against other pan-cancer studies. (A) DEEPEST calls significantly fewer fusions on a subset of GTEx (normal) samples compared to STAR-Fusion used in a recent pan-cancer study. (B) Comparison of the DEEPEST and TumorFusions calls on TCGA normals demonstrates significantly higher specificity for DEEPEST. (C) Number of fusions detected in TCGA tumor samples by DEEPEST and two recent TCGA fusion lists, TCGA fusions and Tumorfusions based on the samples that are processed by all three studies. DEEPEST achieves considerably higher sensitivity on real cancer datasets. (D) Analysis of the fusions called only by one method across tumor types reveals that DEEPEST-only calls are most abundant in cancers known to have high genomic instability such as ESCA and OV.

**Supplemental Figure 4:**
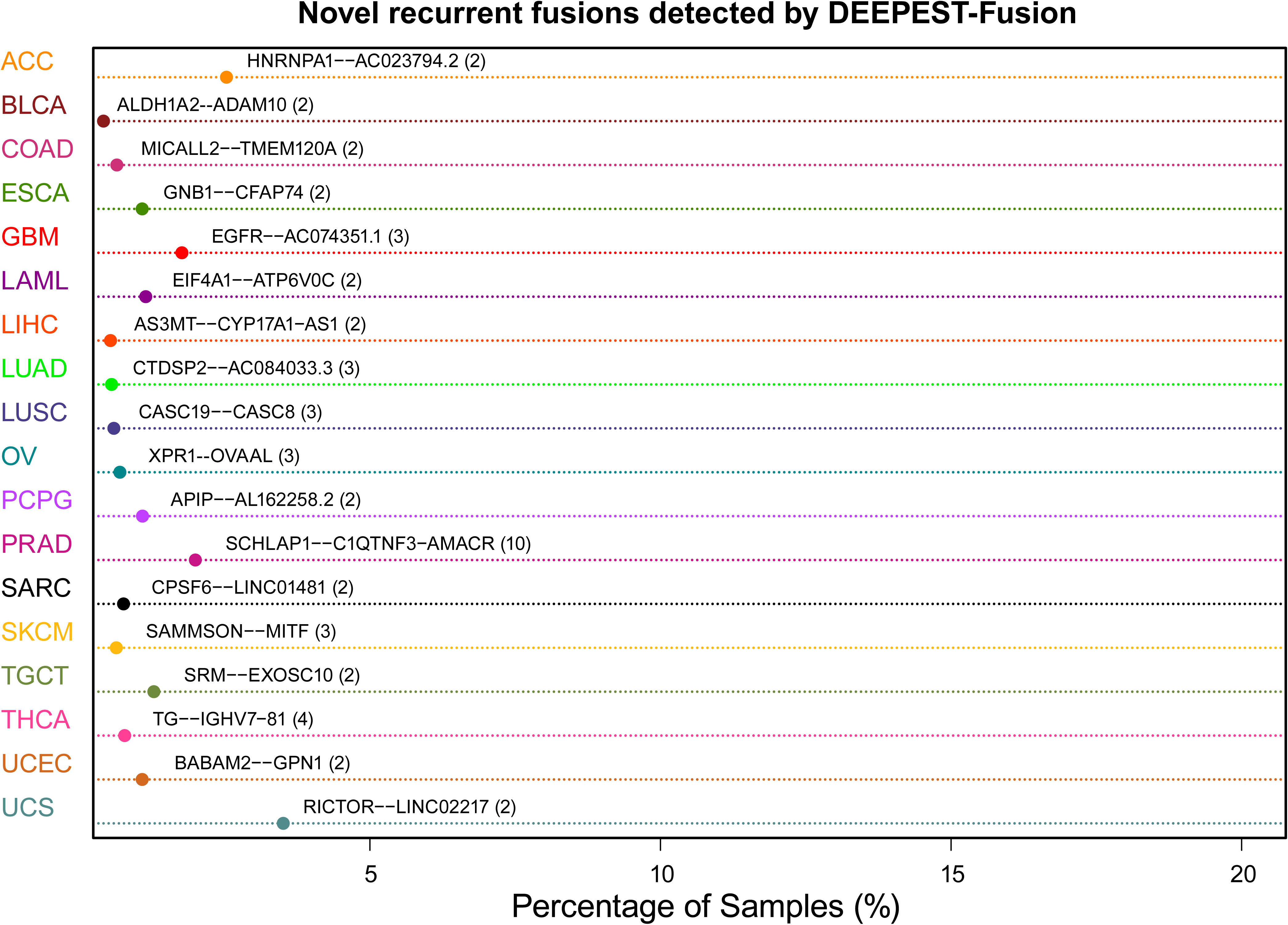
Novel recurrent fusions identified by DEEPEST. DEEPEST calls 157 novel recurrent fusions across various TCGA tumor types. The complete list of novel recurrent fusions is provided in Dataset S1.

**Supplemental Figure 5:**
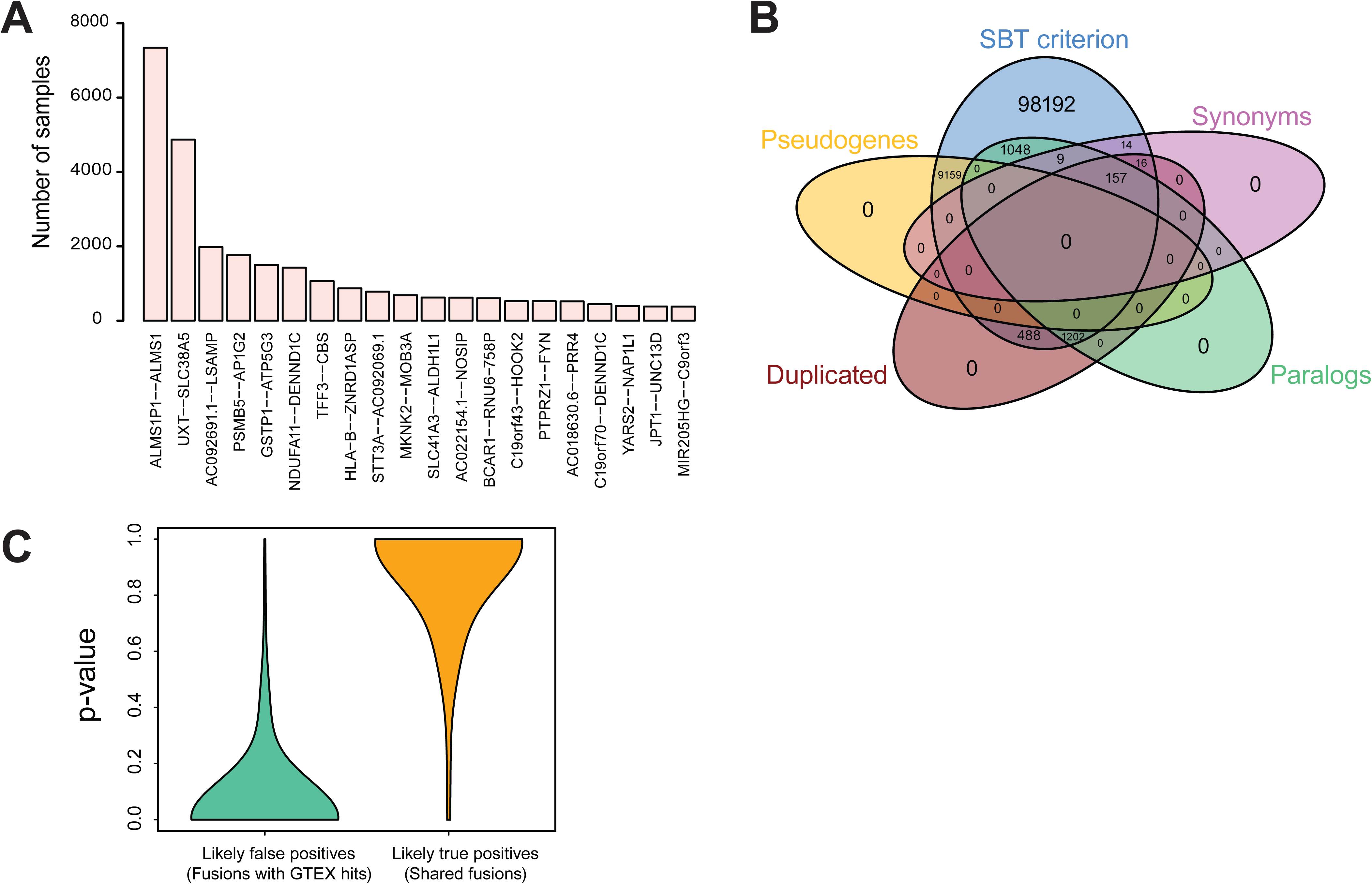
The performance of the SBT-based statistical refinement step. (A) Some of the fusions discarded by the SBT step are filtered out for a large number of samples. (B) A large fraction of the fusions discarded by the statistical refinement step cannot be discarded by other conventional filters based on the biology type of genes. (C) Violin plots show the distribution of the binomial test two-sided p-values between the detection frequency of the DEEPEST first component and that of the SBT querying for fusions nominated by the first component and stratified by likely false positives (fusions with GTEx hits) and likely true positives (shared fusions between the first component of DEEPEST, TumorFusions, and TCGA Fusions). Small p-values indicate that the detection frequencies by the first component of DEEPEST and SBT. This figure demonstrates that while likely true positives will make it through the SBT-based refinement step, likely false positives will be discarded.

**Supplemental Figure 6:**
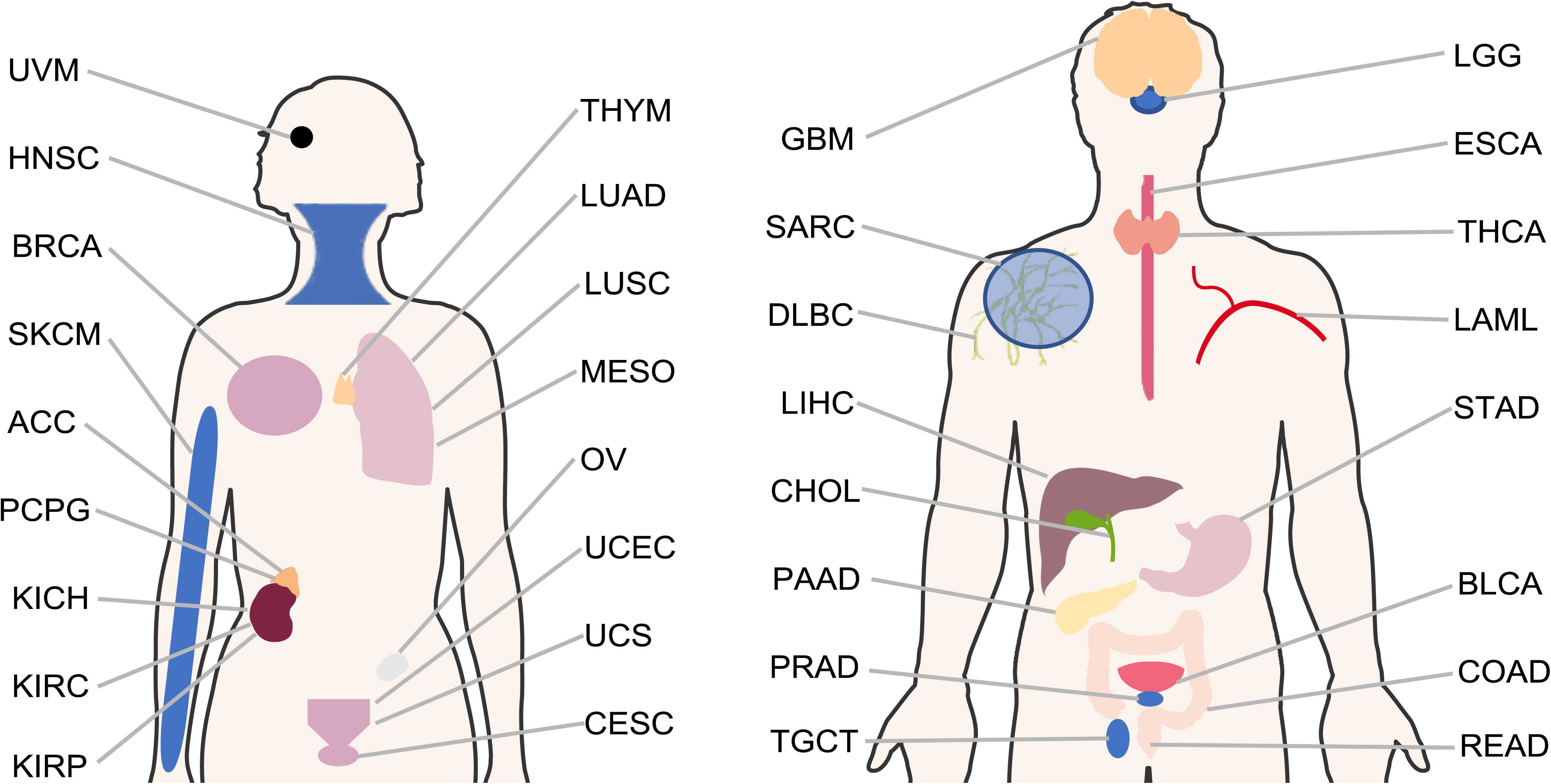
Tumor types analyzed by DEEPEST. Screening all 33 TCGA tumor types by DEEPEST identifies 31,007 high confidence fusions.

**Supplemental Figure 7:**
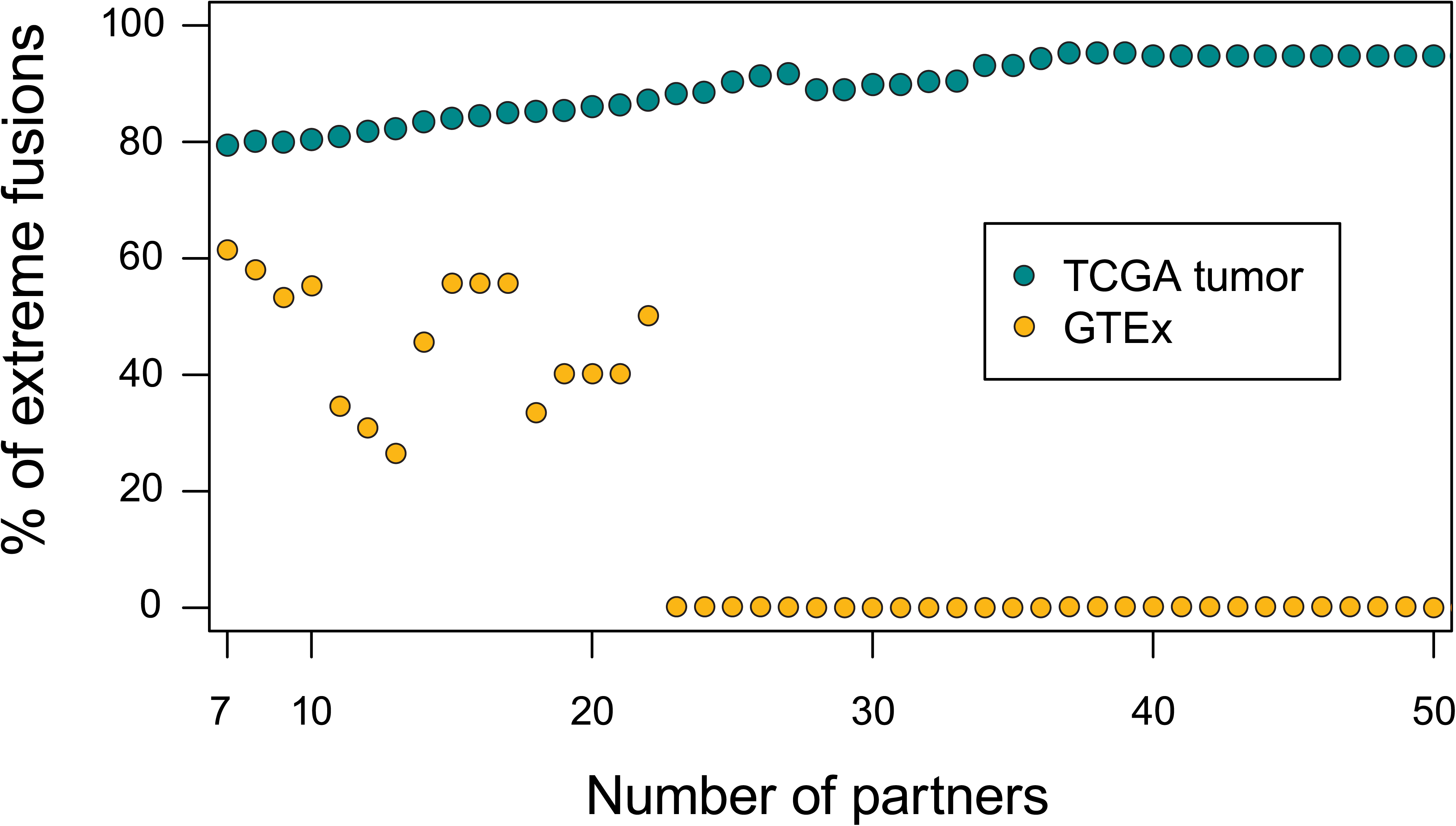
High enrichment of significantly-fused genes in tumors. Fusions with significantly fused genes have different profiles in TCGA (tumor) and GTEx (normal) datasets. While for TCGA tumors, the fraction of extreme fusions increases with the number of distinct partners for significantly fused genes, no extreme fusion involving genes with more than 23 partners is found in GTEx samples.

**Supplemental Figure 8:**
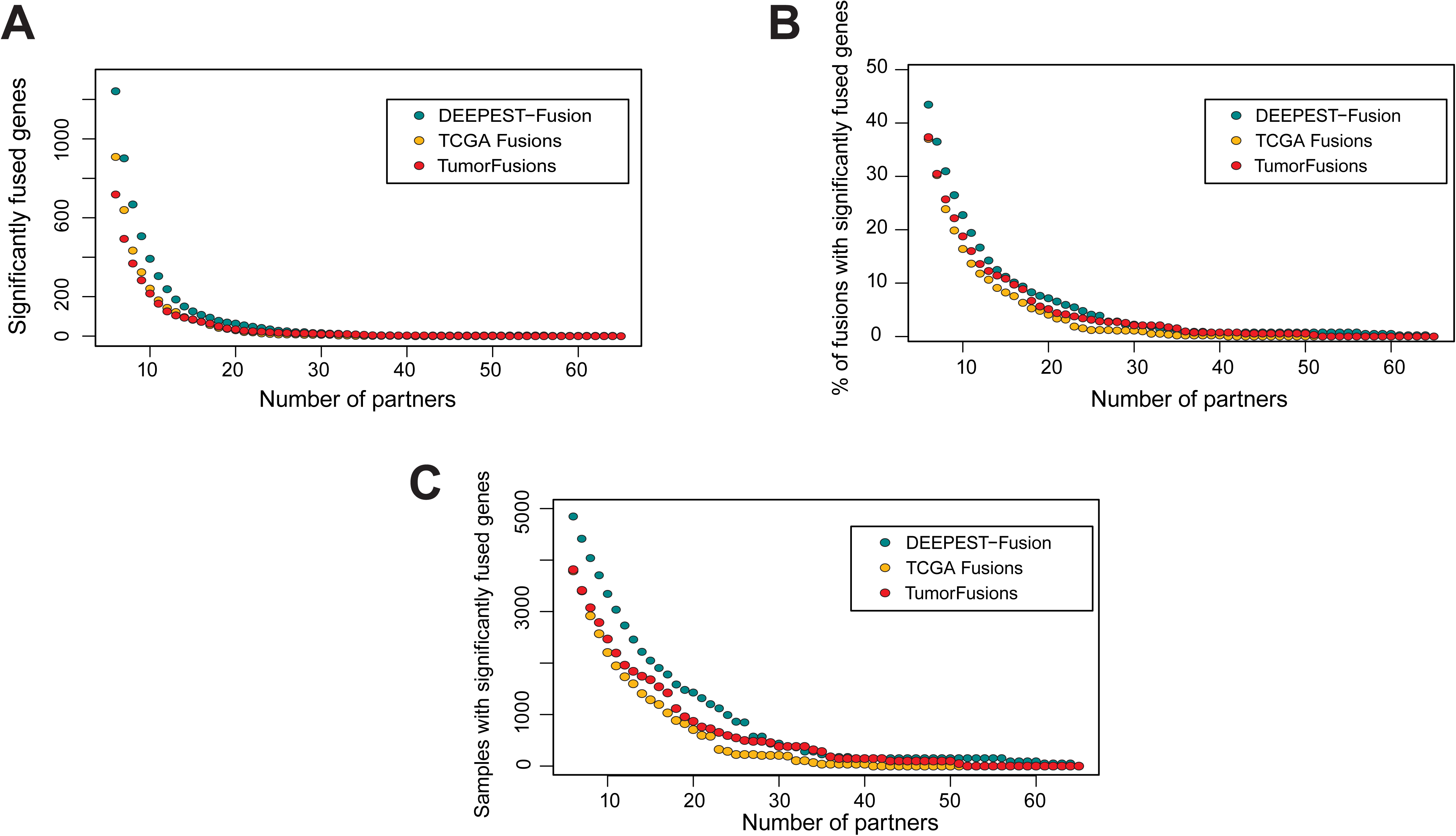
Comparison of the prevalence of significantly-fused genes in recent TCGA studies. (A) DEEPEST calls more significantly fused genes than recent studies. (B) DEEPEST identifies higher enrichment of fusions with significantly fused genes. (C) DEEPEST identifies significantly fused genes in more samples compared to recent studies (similar numbers of samples were analyzed: DEEPEST: 9,946; TumorFusions: 9,966; TCGA Fusions: 9,624).

**Supplemental Figure 9:**
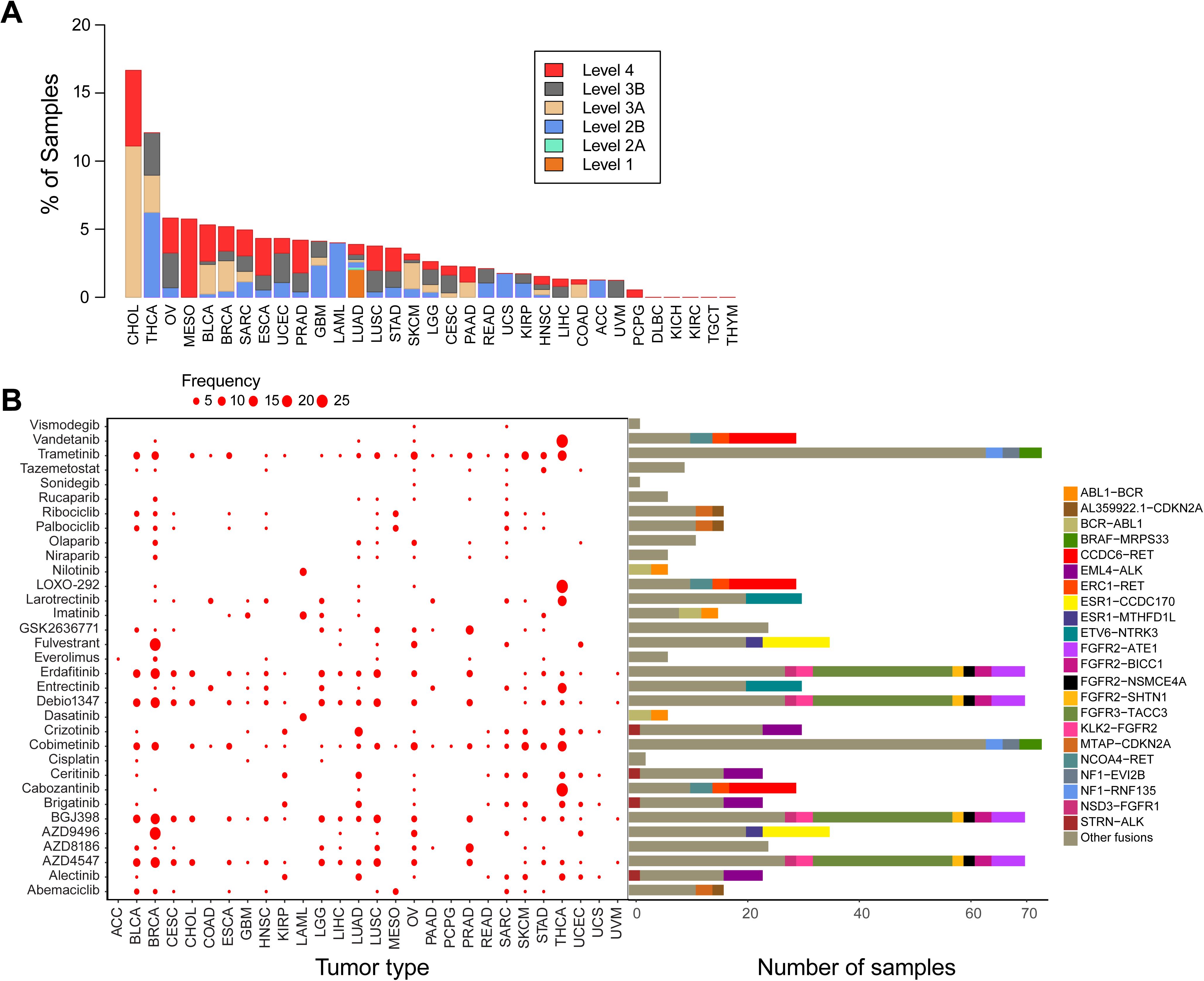
Druggable fusions across various tumor types. (A) Percentage of samples with druggable fusions relative to various levels of drug evidence. Definition of evidence levels: Level 1: FDA-recognized biomarker for an FDA-approved drug; Level 2A: Standard care biomarker for an FDA-approved drug; Level 2B: Standard care biomarker for an FDA approved drug in another indication; Level 3A: biomarker with clinical evidence for a non-standard-care drug; Level 3B: biomarker with clinical evidence for a non-standard-care drug in another indication; Level4: biomarker with biologic evidence for a non-standard-care drug; (B) The dot plot indicates the number of samples with druggable fusion for each drug across various tumor types. The bar chart shows the number of samples involving a certain fusion that can be potentially treated using the given drug.

**Supplemental Figure 10:**
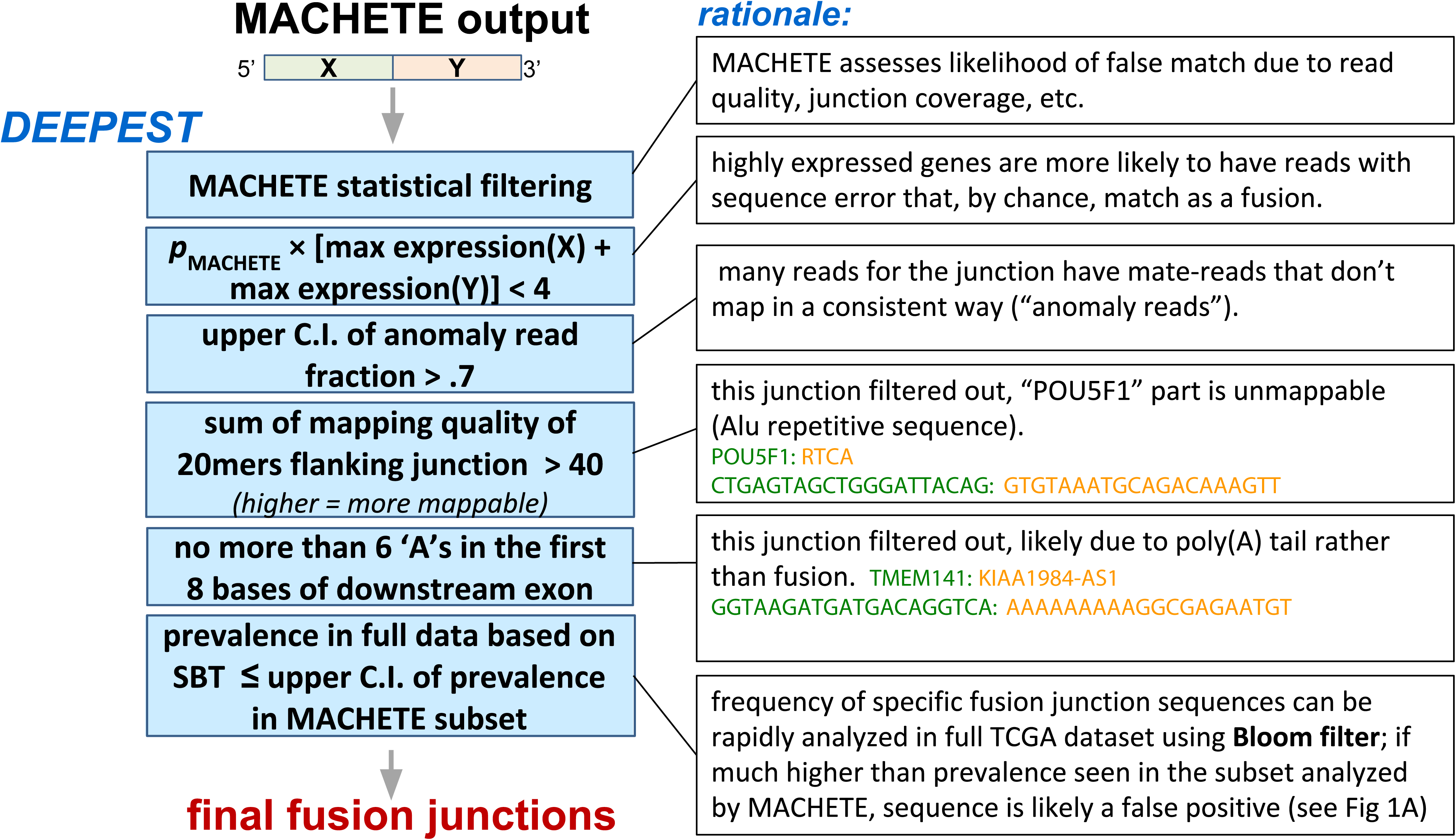
Statistical scores based on the MACHETE modeling are assigned to each junction in junction database. In DEEPEST, junctions that can pass the statistical filtering of MACHETE undergo several additional filters, including p-value correction based on expression values of partner genes, ratio of anomaly reads mapped to the junction, sum of mapping quality values for the 20mers flanking the junction, and the number of A’s in the beginning of downstream exon. A major step in DEEPEST is to check for consistency of detection frequencies obtained by MACHETE and Sequence Bloom Tree (SBT) across the sequencing database, which would mitigate false positive rates due to multiple hypothesis testing in big databases.

## List of Supplemental Tables

Table1: List of fusions called by DEEPEST-Fusion on the TCGA tumor samples, recurrent fusions, novel fusions on TCGA tumors, novel recurrent fusions, detected marker fusions as reported in the TCGA marker papers, and analyzed RNA-Seq files.

Table2: List of significantly fused genes on the 5’ and 3’ sides of fusions, fusions containing at least one significantly fused gene, significantly fused noncoding RNA genes, and GO enrichment analysis for enriched biological processes in significantly fused genes.

Table3: List of protein domains for DEEPEST-Fusion fusions, domain pairs found only in fusion proteins, domain pairs and single domains enriched in fusion proteins, and GO enrichment analysis on enriched protein domains in fusions.

Table4: Fusions containing Kinase amd COSMIC genes and along with the results of the statistical test to find the tumor types that are enriched for kinase and COSMIC fusions.

## References

1. Abate, F., Acquaviva, A., Paciello, G., Foti, C., Ficarra, E., Ferrarini, A., Delledonne, M., Iacobucci, I., Soverini, S., Martinelli, G. and Macii, E. (2012). Bellerophontes: an RNA-Seq data analysis framework for chimeric transcripts discovery based on accurate fusion model. Bioinformatics, 28(16), 2114–2121.

2. Alaei-Mahabadi, B., Bhadury, J., Karlsson, J. W., Nilsson, J. A., & Larsson, E. (2016). Global analysis of somatic structural genomic alterations and their impact on gene expression in diverse human cancers. Proceedings of the National Academy of Sciences, 113(48), 13768–13773.

3. Bailey P, Chang DK, Nones K, Johns AL, Patch AM, Gingras MC, et al. (2016) Genomic analyses identify molecular subtypes of pancreatic cancer. Nature, 531(7592):47–52.

4. Benjamini, Y., & Yekutieli, D. (2001). The control of the false discovery rate in multiple testing under dependency. Annals of Statistics, 1165–1188.

5. Blum, Andrew E., et al. “RNA sequencing identifies transcriptionally viable gene fusions in esophageal adenocarcinomas.” Cancer research 76.19 (2016): 5628–5633.

6. Cai, C., Chen, Q.B., Han, Z.D., Zhang, Y.Q., He, H.C., Chen, J.H., Chen, Y., Yang, S.B., Wu, Y.D., Zeng, Y.R. and Qin, G.Q. (2015). miR-195 inhibits tumor progression by targeting RPS6KB1 in human prostate cancer. Clinical Cancer Research, clincanres- 0217.

7. Cancer Genome Atlas Research Network (2013). Genomic and epigenomic landscapes of adult de novo acute myeloid leukemia. The New England Journal of Medicine, 368(22), 2059–74.

8. Carrara, M., Beccuti, M., Cavallo, F., Donatelli, S., Lazzarato, F., Cordero, F., & Calogero, R. A. (2013). State of art fusion-finder algorithms are suitable to detect transcription-induced fusions in normal tissues? BMC Bioinformatics, 14 Suppl 7(Suppl 7), S2.

9. Cerami, E., Gao, J., Dogrusoz, U., Gross, B.E., Sumer, S.O., Aksoy, B.A., Jacobsen, A., Byrne, C.J., Heuer, M.L., Larsson, E. and Antipin, Y. (2012). The cBio Cancer Genomics Portal: An open platform for exploring multidimensional cancer genomics data. Cancer Discovery, 2(5), 401–404.

10. Chakravarty, D., Gao, J., Phillips, S., Kundra, R., Zhang, H., Wang, J., Rudolph, J.E., Yaeger, R., Soumerai, T., Nissan, M.H. and Chang, M.T. (2017). OncoKB: a precision oncology knowledge base. JCO Precision Oncology, 1, 1–16.

11. Cibulskis, K., Lawrence, M.S., Carter, S.L., Sivachenko, A., Jaffe, D., Sougnez, C., Gabriel, S., Meyerson, M., Lander, E.S. and Getz, G. (2013). Sensitive detection of somatic point mutations in impure and heterogeneous cancer samples. Nature Biotechnology, 31(3), 213–219.

12. Fang, H. (2014). dcGOR: an R package for analysing ontologies and protein domain annotations. PLoS Computational Biology, 10(10), e1003929.

13. Forment, J. V., Kaidi, A., & Jackson, S. P. (2012). Chromothripsis and cancer: causes and consequences of chromosome shattering. Nature Reviews. Cancer, 12(10), 663–70.

14. Gao, Q., Liang, W.W., Foltz, S.M., Mutharasu, G., Jayasinghe, R.G., Cao, S., Liao, W.W., Reynolds, S.M., Wyczalkowski, M.A., Yao, L. and Yu, L. (2018). Driver fusions and their implications in the development and treatment of human cancers. Cell Reports, 23(1), 227–238.

15. Haas, B., Dobin, A., Stransky, N., Li, B., Yang, X., Tickle, T., Bankapur, A., Ganote, C., Doak, T., Pochet, N. and Sun, J. (2017). STAR-Fusion: fast and accurate fusion transcript detection from RNA-Seq. BioRxiv, 120295.

16. Hadari, Y. R., Gotoh, N., Kouhara, H., Lax, I., & Schlessinger, J. (2001). Critical role for the docking-protein FRS2α in FGF receptor-mediated signal transduction pathways. Proceedings of the National Academy of Sciences, 98(15), 8578–8583.

17. Henze, N. (1998). A poisson limit law for a generalized birthday problem. Statistics & Probability Letters, 39(4).

18. Hsieh, G., Bierman, R., Szabo, L., Lee, A.G., Freeman, D.E., Watson, N., Sweet-Cordero, E.A. and Salzman, J. (2017). Statistical algorithms improve accuracy of gene fusion detection. Nucleic Acids Research, 45(13), e126–e126.

19. Hu, X., Wang, Q., Tang, M., Barthel, F., Amin, S., Yoshihara, K., Lang, F.M., Martinez-Ledesma, E., Lee, S.H., Zheng, S. and Verhaak, R.G. (2017). TumorFusions: an integrative resource for cancer-associated transcript fusions. Nucleic Acids Research, 46(D1), D1144–D1149.

20. Huarte, Maite. “The emerging role of lncRNAs in cancer.” Nature medicine 21.11 (2015): 1253.

21. Inaki, K., Hillmer, A.M., Ukil, L., Yao, F., Woo, X.Y., Vardy, L.A., Zawack, K.F.B., Lee, C.W.H., Ariyaratne, P.N., Chan, Y.S. and Desai, K.V. (2011). Transcriptional consequences of genomic structural aberrations in breast cancer. Genome Research, 21(5), 676–687.

22. Kakizuka, A., Miller Jr, W.H., Umesono, K., Warrell Jr, R.P., Frankel, S.R., Murty, V.V.V.S., Dmitrovsky, E. and Evans, R.M. (1991). Chromosomal translocation t (15; 17) in human acute promyelocytic leukemia fuses RARα with a novel putative transcription factor, PML. Cell, 66(4), 663–674.

23. Knudson, A. G. (1971). Mutation and cancer: statistical study of retinoblastoma. Proceedings of the National Academy of Sciences, 68(4), 820–3.

24. Kopp, Florian, and Joshua T. Mendell. “Functional classification and experimental dissection of long noncoding RNAs.” Cell 172.3 (2018): 393–407.

25. Kubbutat, M. H., Jones, S. N., & Vousden, K. H. (1997). Regulation of p53 stability by Mdm2. Nature, 387(6630), 299.

26. Kumar, S., Vo, A. D., Qin, F., & Li, H. (2016). Comparative assessment of methods for the fusion transcripts detection from RNA-Seq data. Scientific Reports, 6, 21597.

27. Latysheva, N. S., & Babu, M. M. (2016). Discovering and understanding oncogenic gene fusions through data intensive computational approaches. Nucleic Acids Research, 44(10), 4487–4503.

28. Lau, J.W., Lehnert, E., Sethi, A., Malhotra, R., Kaushik, G., Onder, Z., Groves-Kirkby, N., Mihajlovic, A., DiGiovanna, J., Srdic, M. and Bajcic, D. (2017). The Cancer Genomics Cloud: collaborative, reproducible, and democratized—a new paradigm in large-scale computational research. Cancer Research, 77(21), e3–e6.

29. Lawrence, M.S., Stojanov, P., Mermel, C.H., Robinson, J.T., Garraway, L.A., Golub, T.R., Meyerson, M., Gabriel, S.B., Lander, E.S. and Getz, G. (2014). Discovery and saturation analysis of cancer genes across 21 tumour types. Nature, 505(7484), 495–501.

30. Lee, M., Lee, K., Yu, N., Jang, I., Choi, I., Kim, P., Jang, Y.E., Kim, B., Kim, S., Lee, B. and Kang, J. (2016). ChimerDB 3.0: an enhanced database for fusion genes from cancer transcriptome and literature data mining. Nucleic Acids Research, 45(D1), D784–D789.

31. Liang, C., Feng, P., Ku, B., Dotan, I., Canaani, D., Oh, B. H., & Jung, J. U. (2006). Autophagic and tumour suppressor activity of a novel Beclin1-binding protein UVRAG. Nature Cell Biology, 8(7), 688.

32. Lim, K.H., Baines, A.T., Fiordalisi, J.J., Shipitsin, M., Feig, L.A., Cox, A.D., Der, C.J. and Counter, C.M. (2005). Activation of RalA is critical for RAS-induced tumorigenesis of human cells. Cancer Cell, 7(6), 533–545.

33. Lin, Chunru, and Liuqing Yang. “Long noncoding RNA in cancer: wiring signaling circuitry.” Trends in cell biology 28.4 (2018): 287–301.

34. Lin, S., Ptasinska, A., Assi, S.A., Kerry, J., Meetei, R.A., Luo, R.T., Thirman, M.J., Milne, T., Bonifer, C. and Mulloy, J.C. (2016). The Transcriptome Heterogeneity of MLL-Fusion ALL Is Driven By Fusion Partners Via Distinct Chromatin Binding. Blood, 128(576).

35. Liu, S., Tsai, W.H., Ding, Y., Chen, R., Fang, Z., Huo, Z., Kim, S., Ma, T., Chang, T.Y., Priedigkeit, N.M. and Lee, A.V. (2015). Comprehensive evaluation of fusion transcript detection algorithms and a meta-caller to combine top performing methods in paired-end RNA-seq data. Nucleic Acids Research, 44(5).

36. Liu, X. S., & Mardis, E. R. (2017). Applications of immunogenomics to cancer. Cell, 168(4), 600–612.

37. Lonsdale, J., Thomas, J., Salvatore, M., Phillips, R., Lo, E., Shad, S., Hasz, R., Walters, G., Garcia, F., Young, N. and Foster, B. (2013). The genotype-tissue expression (GTEx) project. Nature Genetics, 45(6), 580.

38. Lui, G. Y., Grandori, C., & Kemp, C. J. (2018). CDK12: an emerging therapeutic target for cancer. Journal of Clinical Pathology, 71(11), 957–962.

39. Luo, W., Gangwal, K., Sankar, S., Boucher, K. M., Thomas, D., & Lessnick, S. L. (2009). GSTM4 is a microsatellite-containing EWS/FLI target involved in Ewing’s sarcoma oncogenesis and therapeutic resistance. Oncogene, 28(46), 4126.

40. Manning, G., Whyte, D. B., Martinez, R., Hunter, T., & Sudarsanam, S. (2002). The protein kinase complement of the human genome. Science, 298(5600), 1912–1934.

41. Martincorena, I., Roshan, A., Gerstung, M., Ellis, P., Van Loo, P., McLaren, S., Wedge, D.C., Fullam, A., Alexandrov, L.B., Tubio, J.M. and Stebbings, L. (2015). High burden and pervasive positive selection of somatic mutations in normal human skin. Science, 348(6237), 880–886.

42. Minkin, I., Pham, S., & Medvedev, P. (2016). TwoPaCo: An efficient algorithm to build the compacted de Bruijn graph from many complete genomes. Bioinformatics, 33(24), 4024–4032.

43. Negrini, S., Gorgoulis, V. G., & Halazonetis, T. D. (2010). Genomic instability—an evolving hallmark of cancer. Nature Reviews Molecular Cell Biology, 11(3), 220.

44. Nowell, P., & Hungerford, D. (1960). A minute chromosome in human chronic 9 granulocytic leukemia. Science, 132(3438), 1488–1501.

45. Papaemmanuil, E., Gerstung, M., Bullinger, L., Gaidzik, V.I., Paschka, P., Roberts, N.D., Potter, N.E., Heuser, M., Thol, F., Bolli, N. and Gundem, G. (2016). Genomic Classification and Prognosis in Acute Myeloid Leukemia. The New England Journal of Medicine, 374(23), 2202–2221.

46. Prensner, John R., et al. “The long noncoding RNA SChLAP1 promotes aggressive prostate cancer and antagonizes the SWI/SNF complex.” Nature genetics 45.11 (2013): 1392.

47. Ragonnaud E, Holst P. The rationale of vectored gene-fusion vaccines against cancer: evolving strategies and latest evidence. Ther Adv Vaccines. 2013;1(1):33–47.

48. Rowley, J. D. (2001). Chromosome translocations: dangerous liaisons revisited. Nature Reviews Cancer, 1(3), 245.

49. Salzman, J., Gawad, C., Wang, P. L., Lacayo, N., & Brown, P. O. (2012). Circular RNAs are the predominant transcript isoform from hundreds of human genes in diverse cell types. PloS One, 7(2), e30733.

50. Sankar, Savita, and Stephen L. Lessnick. “Promiscuous partnerships in Ewing’s sarcoma.” Cancer genetics 204.7 (2011): 351–365.

51. Saramäki, O.R., Harjula, A.E., Martikainen, P.M., Vessella, R.L., Tammela, T.L., Visakorpi, T. (2008). TMPRSS2: ERG Fusion Identifies a Subgroup of Prostate Cancers with a Favorable Prognosis. Clinical Cancer Research, 14(11), 3395–400.

52. Singh, D., Chan, J.M., Zoppoli, P., Niola, F., Sullivan, R., Castano, A., Liu, E.M., Reichel, J., Porrati, P., Pellegatta, S. and Qiu, K. (2012). Transforming fusions of FGFR and TACC genes in human glioblastoma. Science, 337(6099), 1231–1235.

53. Soda, M., Choi, Y.L., Enomoto, M., Takada, S., Yamashita, Y., Ishikawa, S., Fujiwara, S.I., Watanabe, H., Kurashina, K., Hatanaka, H. and Bando, M. (2007). Identification of the transforming EML4-ALK fusion gene in non-small-cell lung cancer. Nature, 448(7153), 561–566.

54. Solomon, B. and Kingsford, C. (2016). Fast search of thousands of short-read sequencing experiments. Nature Biotechnology, 34(3): 300–302.

55. Stranger, B. E., Forrest, M. S., Dunning, M., Ingle, C. E., Beazley, C., Thorne, N., … & Tyler-Smith, C. (2007). Relative impact of nucleotide and copy number variation on gene expression phenotypes. Science, 315(5813), 848–853.

56. Stransky, N., Cerami, E., Schalm, S., Kim, J. L., & Lengauer, C. (2014). The landscape of kinase fusions in cancer. Nature Communications, 5, 4846.

57. Szabo, L., Morey, R., Palpant, N.J., Wang, P.L., Afari, N., Jiang, C., Parast, M.M., Murry, C.E., Laurent, L.C. and Salzman, J. (2015). Statistically based splicing detection reveals neural enrichment and tissue-specific induction of circular RNA during human fetal development. Genome Biology, 16, 126.

58. Thacker, J. (2005). The RAD51 gene family, genetic instability and cancer. Cancer Letters, 219(2), 125–135.

59. Tomlins, S.A., Laxman, B., Varambally, S., Cao, X., Yu, J., Helgeson, B.E., Cao, Q., Prensner, J.R., Rubin, M.A., Shah, R.B. and Mehra, R., (2008). Role of the TMPRSS2-ERG gene fusion in prostate cancer. Neoplasia, 10(2), IN1–IN9.

60. Torres-García, W., Zheng, S., Sivachenko, A., Vegesna, R., Wang, Q., Yao, R., Berger, M.F., Weinstein, J.N., Getz, G. and Verhaak, R.G. (2014). PRADA: pipeline for RNA sequencing data analysis. Bioinformatics, 30(15), 2224–2226.

61. Yoshihara, K., Wang, Q., Torres-Garcia, W., Zheng, S., Vegesna, R., Kim, H., & Verhaak, R. G. W. (2015). The landscape and therapeutic relevance of cancer-associated transcript fusions. Oncogene, 34(37), 4845–54.

62. Zhang J, Mardis ER, Maher CA. INTEGRATE-neo: a pipeline for personalized gene fusion neoantigen discovery. Bioinformatics. 2017;33(4):555–7.

63. Sun, C. C., Li, S. J., Li, G., Hua, R. X., Zhou, X. H., & Li, D. J. (2016). Long Intergenic Noncoding RNA 00511 acts as an oncogene in non–small-cell lung cancer by binding to EZH2 and suppressing p57. Molecular Therapy-Nucleic Acids, 5, e385.

64. Xing, Z., Lin, A., Li, C., Liang, K., Wang, S., Liu, Y., … & Hung, M. C. (2014). lncRNA directs cooperative epigenetic regulation downstream of chemokine signals. Cell, 159(5), 1110–1125.

65. Guan, Y., Kuo, W. L., Stilwell, J. L., Takano, H., Lapuk, A. V., Fridlyand, J., … & Kalloger, S. E. (2007). Amplification of PVT1 contributes to the pathophysiology of ovarian and breast cancer. Clinical cancer research, 13(19), 5745–5755.

